# Comprehensive Transcriptomic Analysis of Atopic Dermatitis Patients Documents the Spectrum of Molecular Abnormalities and the Response to Treatment

**DOI:** 10.64898/2026.03.04.709610

**Authors:** Sneha Shrotri, Andrea Daamen, Prathyusha Bachali, Amrie Grammer, Peter E Lipsky

**Author notes:** **Correspondence:** Peter E Lipsky.

## Abstract

Atopic dermatitis (AD) is a chronic inflammatory skin disease characterized by immune dysregulation and barrier dysfunction. To define the molecular architecture of AD in greater detail, we integrated lesional (LES) and non-lesional (NLS) transcriptomic data from multiple datasets using gene expression data from normal skin and psoriasis (PSO) and nummular eczema (NME) cohorts as reference. Gene set variation analysis revealed that adult AD exhibits broad immune activation and consistent barrier impairment in both LES and NLS skin, whereas pediatric AD is dominated by IL-1–driven inflammation with minimal barrier alteration. Comparative analyses showed stronger Th2 and myeloid activity in AD, metabolic enrichment in PSO, and complement and NK cell activation in NME. Longitudinal profiling identified temporal variation in Th1, Th2 and IFN pathways in AD skin. An eczema immune and cellular score, ECZECIS, was developed to quantify transcriptomic abnormalities and correlated with clinical improvement following dupilumab therapy. Among all treatments analyzed, dupilumab produced the most extensive reduction of immune and cytokine pathway activity in skin and attenuated systemic immune activation in blood. These findings delineate distinct immune and barrier signatures across age groups and disease types and establish ECZECIS as a quantitative biomarker for monitoring molecular treatment response in AD.

## INTRODUCTION

Atopic dermatitis (AD) is a chronic, relapsing inflammatory skin disorder that affects both children and adults, with significant heterogeneity in its clinical presentation, underlying immunopathology and treatment response (Bieber 2008; Brunner et al. 2017a; Fishbein et al. 2020; Jeskey et al. 2024, Lobefaro et al. 2022; Torres et al. 2025). Although recent advances in targeted therapies have transformed the treatment landscape (Ratchataswan et al. 2021), the molecular mechanisms driving disease heterogeneity across age groups, disease states, and therapeutic interventions remain incompletely understood (Nomura et al. 2020; Sroka-Tomaszewska and Trzeciak 2021).

Transcriptomic profiling has provided an unprecedented view into the cellular and molecular networks underlying AD, revealing abnormalities in immune activation, cytokine signaling, and epidermal barrier function (Wu et al. 2021). However, existing studies have been limited in scope, often focusing on a single cohort, age group or treatment, and few have systematically compared AD with other inflammatory skin conditions (Seremet et al. 2024) or integrated longitudinal and therapeutic datasets (Fukushima-Nomura et al. 2025). A deeper understanding of these molecular mechanisms is necessary to understand disease pathogenesis, define shared versus unique pathways across skin disorders and identify robust biomarkers for treatment more completely.

In addition to skin pathology, recent blood-based transcriptomic studies have demonstrated that AD is associated with systemic immune dysregulation (Brunner et al. 2017b; Hubbard et al. 2023; Möbus et al. 2022). Incorporating blood transcriptomes alongside skin analysis provides a more comprehensive picture of disease pathogenesis and offers opportunities to identify accessible biomarkers for treatment monitoring.

In this study, we carried out a comprehensive analysis of publicly available datasets to examine gene expression changes across lesional (LES), non-lesional (NLS) and healthy control (CTL) skin in both adult and pediatric AD patients. Using a curated panel of 56 immune, metabolic, tissue-specific and skin barrier-related gene signatures (Martínez et al. 2022), we characterized molecular abnormalities in skin of adult AD patients and investigated differences with pediatric AD and other inflammatory skin diseases, temporal dynamics of AD skin over the course of disease and the impact of different therapeutic interventions on transcriptomic abnormalities. To validate these analyses, we also applied machine-learning models and demonstrated an accuracy of >90% in predicting LES AD skin and also identified the signatures that contributed most strongly to classification accuracy based on Gini importance. To quantify overall molecular alterations, we further developed a composite transcriptomic score, ECZECIS (Eczema Cell and Immune Score), and assessed its association with clinical severity and treatment response. Finally, we extended our analysis to blood transcriptomes to evaluate systemic molecular changes and determine how baseline immune endotypes influence therapeutic outcomes.

Together, these analyses provide an integrated framework for understanding the molecular drivers of AD across patient populations, disease states and therapeutic contexts. The results highlight both shared and distinct pathways that contribute to AD pathogenesis, underscore differences between adult and pediatric disease and establish ECZECIS as a potential molecular biomarker of treatment response.

## RESULTS

### Gene expression profiling reveals molecular differences related to immune and barrier functions in adult and pediatric AD patients

To explore the molecular mechanisms underlying AD pathogenesis, we compared baseline gene expression profiles from five publicly available datasets (Table S1) focusing on both LES and NLS skin of adult and pediatric AD patients in relation to CTL. We conducted Gene Set Variation Analysis (GSVA) using 56 curated gene signatures (Martínez et al. 2022; Shrotri et al. 2024; Table S2), grouped into seven functional categories: immune cells, tissue cells, immune cell processes, cytokines, metabolism, T cell subsets and skin barrier function. Statistical differences in GSVA scores were visualized using Hedge’s g effect-size heatmaps and violin plots (Figure 1a; Figure S2). Patient-level GSVA heatmaps across all datasets further revealed distinct expression patterns in LES, NLS and CTL skin for both adult and pediatric cohorts (Figure S1a-b).

**Figure 1:**
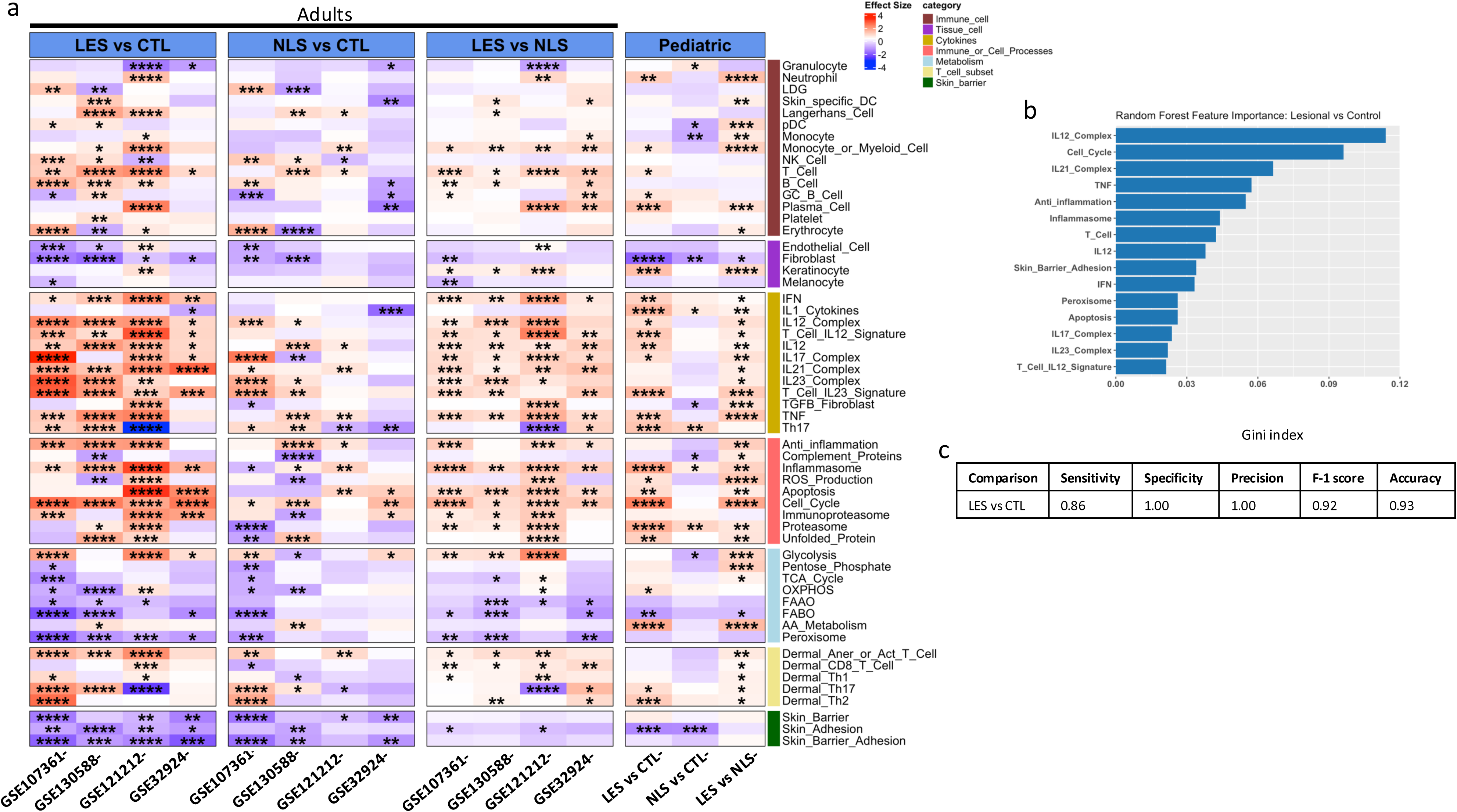
Molecular distinctions between LES, NLS, and CTL skin in atopic dermatitis at baseline: **a.** The heatmap depicts magnitude of change in GSVA scores for each gene signature expressed as Hedge’s G effect sizes, across different comparisons: baseline LES vs NLS skin, and both LES and NLS compared to CTL. Red represents an increase in effect size, while blue denotes a decrease. Asterisks indicate statistical significance, determined using Welch’s t-test for unpaired samples or paired t-test for paired samples. Significance levels are denoted by: *P < 0.05, **P < 0.01, ***P < 0.001, and ****P < 0.0001. **b**. Horizontal bar plot showing the top GSVA-derived molecular signatures contributing to classification of LES vs CTL skin using a Random Forest model. **c.** Summary table reporting Random Forest classifier performance metrics.

In adult AD LES skin, considerable heterogeneity was observed across datasets; therefore, we defined consensus abnormalities as gene signatures enriched in at least three of the four datasets. Using this criterion, several immune-related signatures were consistently upregulated in adult LES skin compared to CTL, including B cells, anergic or activated T cells, IL-21 and IL-23 cytokine complexes, anti-inflammatory pathways and the immunoproteasome (Figure 1a; Figure S1c). In addition, skin barrier function signatures were uniformly downregulated across all adult datasets, underscoring the well-established role of barrier impairment in AD (Cork et al. 2009; Elias et al. 2008; Weidinger and Novak 2016). To validate these findings, we applied various supervised machine-learning models on GSVA scores of LES and CTL skin (Figure S3). As a result, the random forest (RF) classifier, for example, achieved high performance (Sensitivity = 0.86, Specificity = 1.00, Precision = 1.00, F1 = 0.92, Accuracy = 0.93), confirming that GSVA-based pathway scores accurately discriminate LES AD from CTL skin (Figure 1c). Feature-importance analysis based on the Gini index identified IL-12 complex, cell cycle pathways, IL-21 complex, TNF signaling, anti-inflammatory responses and inflammasome activation as the top drivers of LES skin classification (Figure 1b). These ML-derived features closely aligned with biological abnormalities observed in the heatmaps and effect-size analyses, providing orthogonal validation of the key molecular pathways distinguishing LES AD skin from CTL. Adult NL AD skin also showed broad immune activation, with significant upregulation of Langerhans cell and T cell signatures, IL-12 and IL-23 cytokine complexes, TNF signaling, anti-inflammatory pathways, Th17 responses and the cell cycle module. Importantly, adult NL AD skin also displayed downregulation of barrier function signatures (Figure 1a).

In comparison, the immune pathways upregulated in adults were largely absent in pediatric AD patients, whereas the IL-1 cytokine signature was specifically enriched in pediatric LES skin but not in adults (Figure S1c). Furthermore, pediatric LES skin did not show significant alterations in barrier-related pathways, suggesting that barrier dysfunction may be less pronounced in early-onset AD. Pediatric NLS AD skin was enriched for granulocyte, proteasome, and IL-1 cytokine–related pathways, whereas plasmacytoid dendritic cell (pDC) and monocyte signatures were significantly downregulated. In addition, similar to pediatric LES AD skin, skin barrier function gene signatures were not changed in pediatric NLS AD skin. In order to determine whether these results were influenced by age-dependent transcriptional differences, we directly compared healthy adult and healthy pediatric skin biopsies (Figure S4). Among the gene signatures that were differentially enriched in adult as compared to pediatric AD, many, including IL-1 cytokines, were heterogeneous across both adult and pediatric skin, whereas others including the skin barrier and skin adhesion signatures were notably decreased in healthy children. Taken together, these findings emphasize both the shared and age-specific molecular alterations in AD.

### Transcriptomic analysis demonstrates immune and metabolic differences between AD, PSO and NME patients

To delineate the molecular features that are unique to AD and those that are shared with other inflammatory skin diseases, we carried out a comparative analysis of LES and NLS skin from AD with psoriasis (PSO) and nummular eczema (NME), two clinically distinct but pathologically overlapping skin disorders. AD LES skin showed dysregulation of numerous immune and inflammatory pathways. Signatures associated with monocytes and T cell IL-12 responses were significantly upregulated, whereas pathways linked to Langerhans cells, peroxisomal function, fatty acid β-oxidation (FABO) and skin barrier/adhesion were significantly downregulated (Figure 2). Notably, the magnitude of these transcriptomic changes was greater in AD than in PSO (Figure S5).

**Figure 2:**
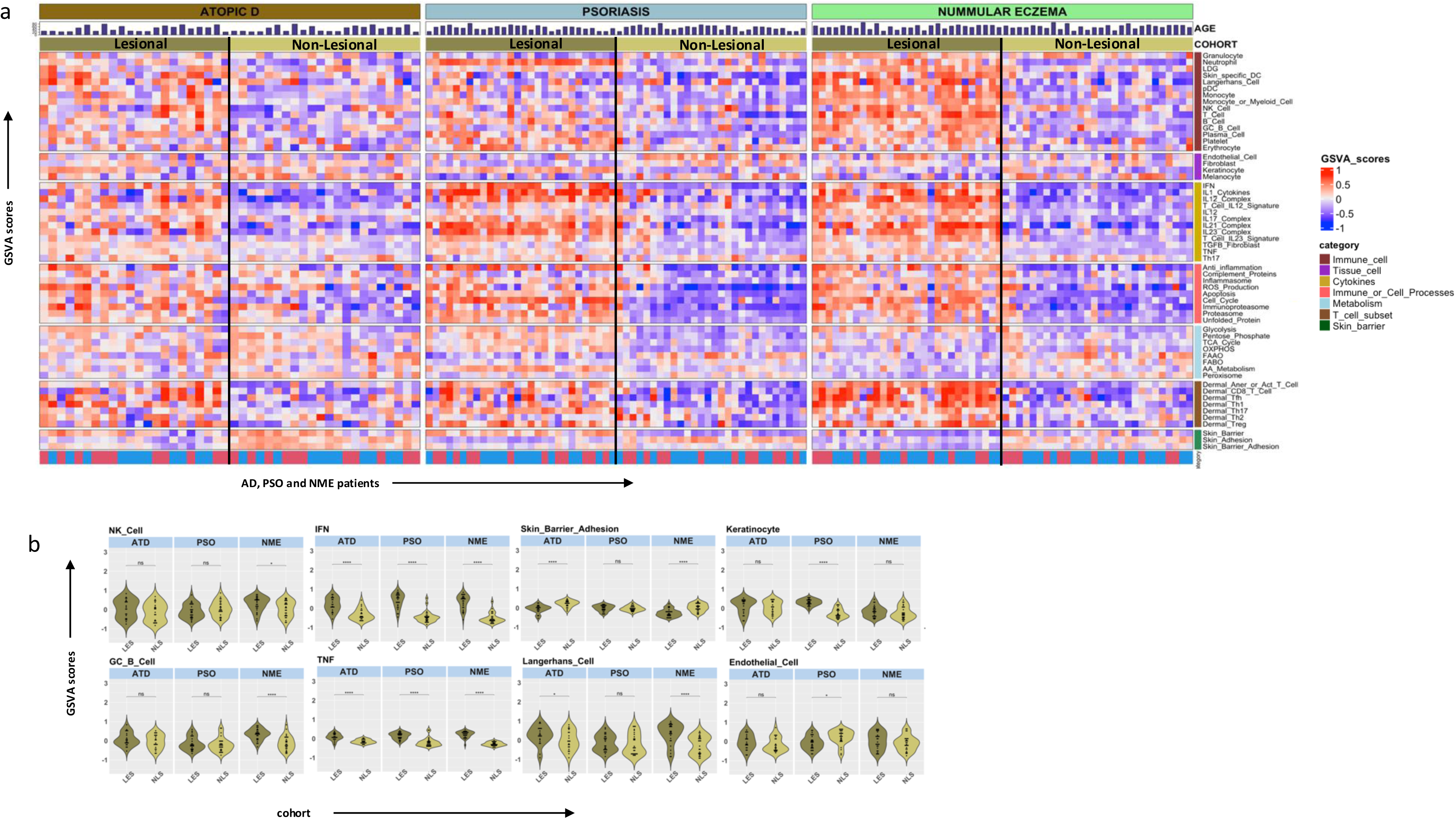
Comparison of LS and NL transcriptomic profiles of AD, psoriasis, and nummular eczema patients: **a**. Heatmap displaying GSVA scores of LS and NL skin of AD, psoriasis, and nummular eczema patients. **b**. Violin plots of NK cell, GC B cell, IFN, TNF, Skin barrier adhesion, Langerhans cell, Keratinocytes, and Endothelial cell signatures showing statistical differences between LS and NL skin for each disease. Dots signify individual patient GSVA scores for each disease and violin silhouettes signify the distribution of patient scores. *P < 0.05; **P < 0.01; ***P < 0.001; ****P < 0.0001

By contrast, PSO was characterized by a distinct cellular and metabolic profile, including enrichment of keratinocyte, pentose phosphate pathway, tricarboxylic acid (TCA) cycle, oxidative phosphorylation (OXPHOS) and amino acid metabolism signatures (p < 0.0001 for all) (Figure 2; Figure S5). Interestingly, the skin barrier signature was upregulated in PSO, but downregulated in both AD and NME. NME patients, meanwhile, exhibited unique immune features, such as enrichment of natural killer (NK) cell, germinal center (GC) B cell and complement protein signatures, suggesting a distinct immune response not shared with AD or PSO. (Figure 2, Figure S5). Overall, LES skin from all three inflammatory conditions demonstrated significant enrichment of immune and cytokine signatures compared with NLS skin. However, the magnitude of enrichment varied across diseases, with NME patients showing the strongest immune activation. (Figure S5)

### Temporal dynamics of immune gene signatures in LES and NLS skin of AD patients

Next, we examined the stability of gene expression profiles by analyzing skin biopsies of AD patients and CTLs taken at multiple timepoints (baseline, month 3, 6, 9 and 12) over the course of one year (Figure S6-7). The individual patient fluctuations in GSVA scores of select gene signatures over time are visualized as line graphs (Figure 3). In CTL samples, gene expression profiles were generally stable, and did not follow any observable patterns across individuals over a year of sampling (Figure 3). Notably, the only significant increase in enrichment was observed in the inflammasome gene signature between baseline and month 3 (p = 0.027) (Figure S7; Table S3). In the LES skin of AD patients, certain pathways showed common fluctuations across individuals, including the IL12 signature, which was increased across patients, and the skin barrier signature, which was generally decreased across patients over time (Figure 3). In addition, comparison of individual time points revealed significant changes in gene signature enrichment (Figure S7, Table S3). The Th1 gene signature showed a significant increase from months 3 to 6 (p = 0.03) indicating an enrichment of Th1-related immune responses in this period. The Th2 gene signature, commonly associated with AD pathology (Gittler et al. 2012; Noda et al. 2015), was significantly upregulated from baseline to month 6 (p = 0.04). Interferon (IFN)-responsive genes were significantly decreased between baseline and month 9 (p = 0.046), whereas IL12 signaling showed a significant increase between baseline and month 12 (p = 0.025). These results suggest a more dynamic immune response in LES skin of AD patients compared to CTL, with Th1 and Th2 pathways playing distinct roles at different stages of the disease progression (Figure S7, Table S3).

**Figure 3:**
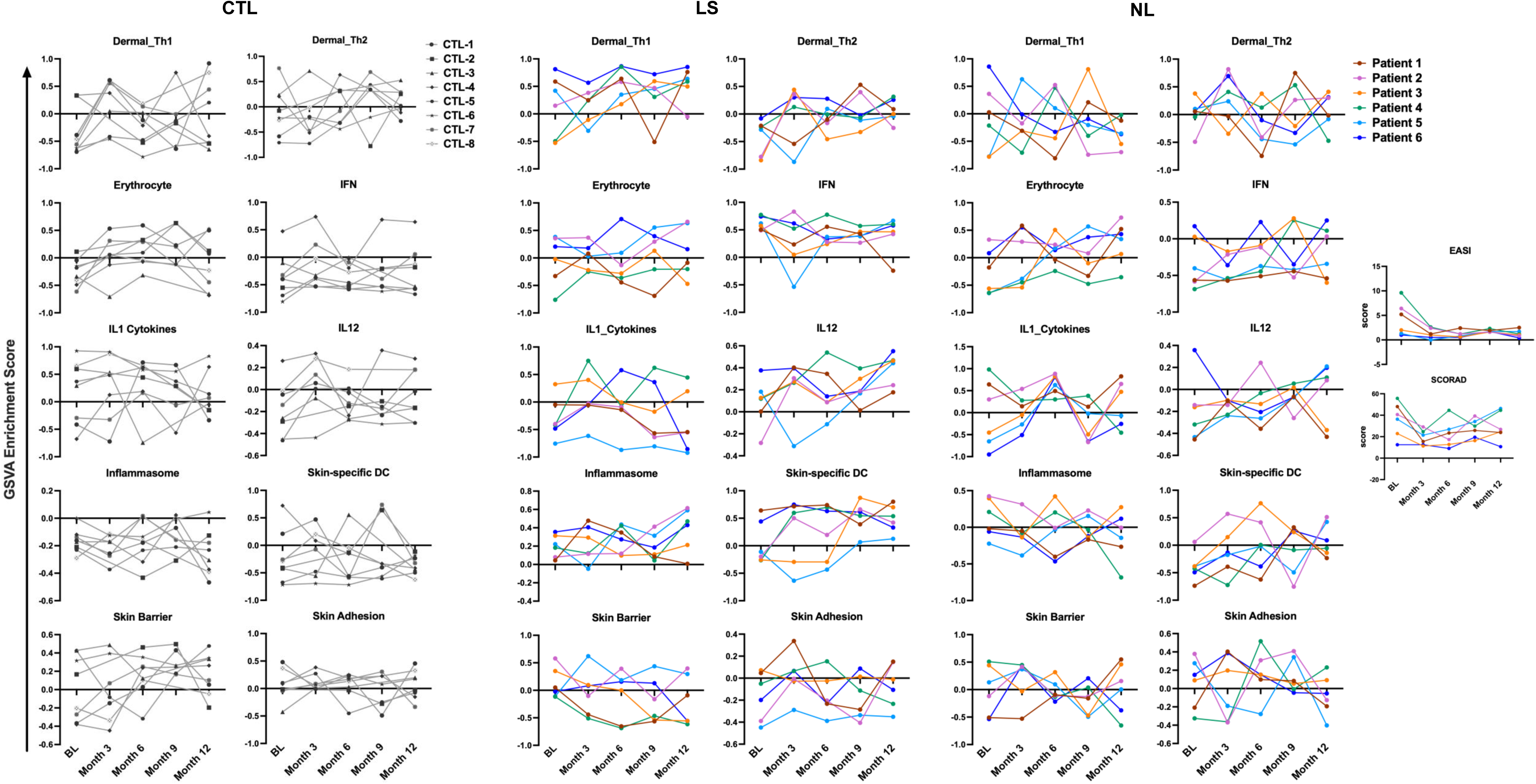
Dynamic changes in AD pathogenesis over time: Line graphs depicting longitudinal GSVA, EASI, and SCORAD scores at baseline (BL), month 3, month 6, month 9, and month 12 for CTL, LS, and NL skin biopsies of AD patients. Each CTL score is denoted by symbol, and each AD patient is denoted by line color.

The longitudinal gene signature enrichment profile of NLS skin of AD patients resembled CTL skin to a greater extent than LES skin, in particular in relation to inflammatory cytokine signatures including IFN and IL12 pathways, which were largely negatively enriched over time (Figure 3; Figure S6-7). Interestingly, the IL1 cytokine gene signature showed a uniform pattern over time with peaks at 6 months and 12 months across patients. Significant changes in GSVA scores of NLS skin were only observed in erythrocyte and skin-specific dendritic cell (DC) gene signatures between baseline and month 12 (p=0.036, p=0.015) (Table S3). It is notable that clinical measures of skin involvement, including the EASI and SCORAD generally decreased from baseline to month 3 and remained stable for the rest of the time course (Figure 3; Figure S6-7). However, there was no correlation between the clinical assessments and any of the gene modules that were variable in either LES or NLS skin. In summary, these results highlight the distinct temporal patterns of immune activation in AD, particularly the variation in Th1, Th2, and IFN pathways in LES skin. Whether these changes reflect responses to standard of care medication or natural variation in gene expression is not known, but these variations do not appear to correlate with clinical activity.

### Assessing molecular abnormalities with the Eczema Cell and Immune Score (ECZECIS)

To summate molecular abnormalities in LES skin of AD patients and incorporate changes in the entire gene expression profile, we developed a transcriptomic scoring system, ECZECIS. To derive ECZECIS, we first carried out GSVA independently on each of four datasets using baseline LES and CTL samples. GSVA scores were then combined across datasets, and ridge-penalized logistic regression was applied to distinguish LES from CTL samples. The resulting regression coefficients were then used to compute ECZECIS for each patient (Figure S8, see Methods). At baseline in the above mentioned longitudinal dataset of untreated AD patients, ECZECIS scores varied widely in the patients, and over time there was modest but non-significant variation (Table S4). To determine whether there was an association between ECZECIS and clinical disease severity, as measured by the EASI and SCORAD, we carried out Pearson’s correlation analysis and found a weak and statistically non-significant correlation between ECZECIS and the EASI score (R = 0.05, p = 0.79) or SCORAD (R=0.12, p=0.54) (Figure 4a), This result confirmed that the molecular alterations captured by the transcriptomic score may not directly reflect changes in clinical disease severity in patients treated with standard of care medication. To further assess how molecular changes relate to clinical outcomes, we focused on the dupilumab treatment dataset (GSE130588), that included matched EASI and SCORAD scores for placebo- and dupilumab-treated patients at baseline, week 4 and week 16 (Figure 4c-d; Table S1). In placebo treated patients, there were no significant changes in ECZECIS in LES skin or clinical severity scores, whereas significant decreases in ECZECIS and clinical severity scores were noted within 4 weeks of treatment with dupilumab. Next, we evaluated the relationship between ECZECIS in LES skin and clinical severity scores. In the placebo group, ECZECIS showed a weak-to-moderate correlation with EASI (R = 0.44, p = 0.07) and SCORAD (R = 0.50, p = 0.03). In contrast, in the dupilumab group there were significant correlations between ECZECIS and clinical scores (EASI: R = 0.46, p = 0.003; SCORAD: R = 0.58, p < 0.0001), suggesting that ECZECIS effectively captures molecular improvements that align with clinical response (Figure 4c). To determine the magnitude of response to dupilumab, we compared ECZECIS scores in LES and NLS skin of treated patients. At baseline ECZECIS scores were significantly greater in LES compared to NLS skin, whereas after treatment (Figure 4d), ECZECIS scores were similar, suggesting that dupilumab had decreased dermal inflammation to NLS levels, but not to that of CTL skin. These findings underscore the utility of transcriptomic profiling, particularly ECZECIS, as a molecular biomarker of therapeutic response in AD, complementing traditional clinical outcome measures.

**Figure 4:**
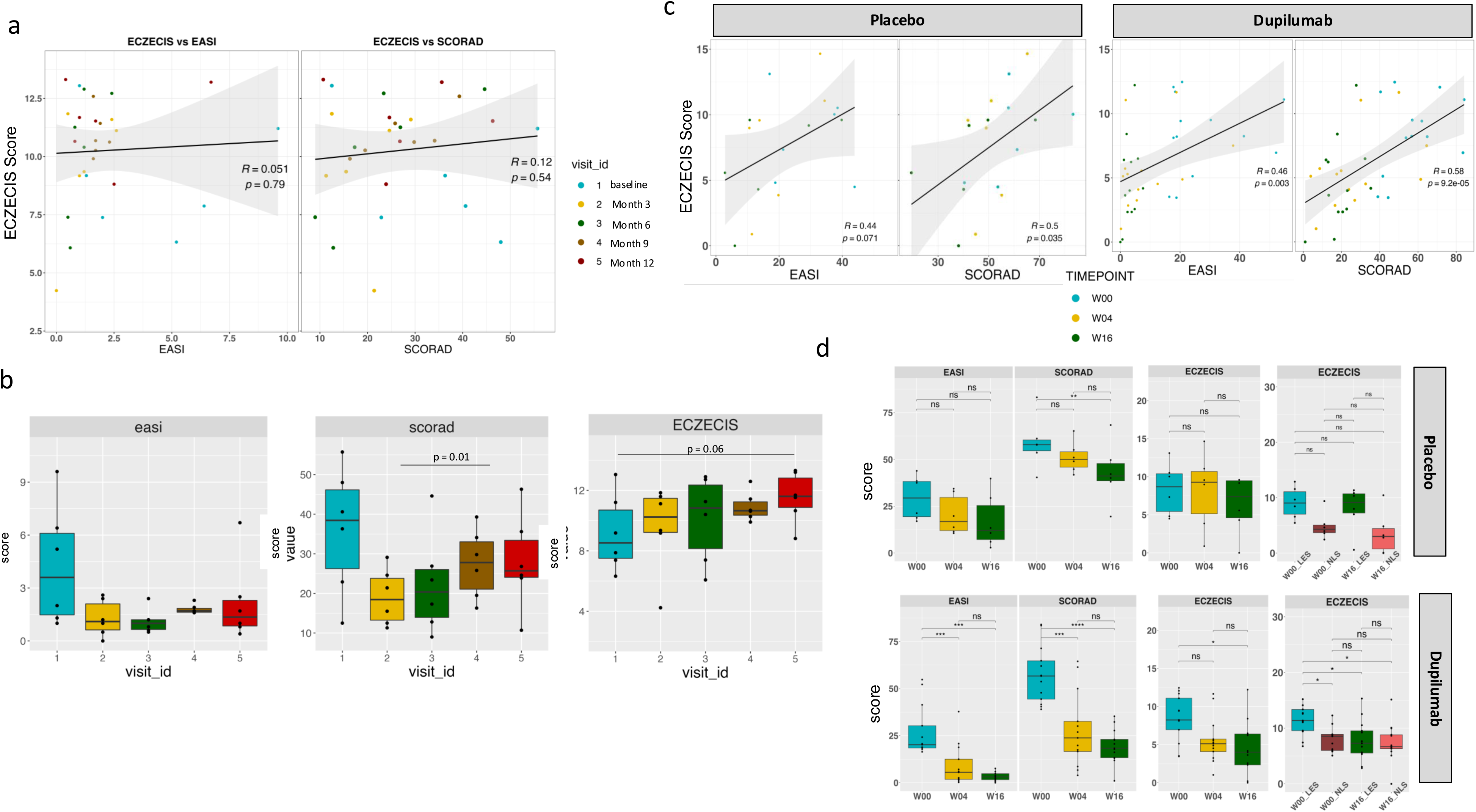
Derivation and correlation of Eczema Cell and Immune Score (ECZECIS): **a-b** Correlation plots depicting the Pearson correlation between ECZECIS and clinical measures of disease severity, EASI and SCORAD a. GSE19903 dataset b. GSE130588 dataset (placebo and dupilumab separately) **c-d**: Boxplots showing changes in EASI, SCORAD, and ECZECIS scores over time, with statistical significance determined using paired t-tests and bonferroni-adjusted p-values.

### Longitudinal changes in gene expression profiles following therapeutic intervention in AD patients

To examine treatment responses, we analyzed transcriptomic profiles from LES biopsies of AD patients treated with a range of therapeutic agents, including placebo/vehicle, cyclosporine, dupilumab, fezakinumab, crisaborole, cedulatinib and apremilast (30 mg and 40 mg) (Table S1). Sampling time points varied by study: placebo and dupilumab (GSE130588) at baseline, week 4, and week 16; cyclosporine or dupilumab (GSE157194) at baseline and month 3; fezakinumab (GSE99802) at baseline, week 4, and week 12; crisaborole (GSE133477) at baseline, day 8 and day 15; cedulatinib (GSE141570) at baseline and day 14; and apremilast (GSE120899) at baseline and week 12. Because of the non-overlapping collection points across studies, we evaluated longitudinal changes within each treatment arm by GSVA using the curated panel of immune, skin barrier, metabolic and tissue-specific gene signatures.

Among all agents, the IL4/IL13 targeting agent dupilumab induced the most extensive transcriptomic alterations by week 16 of therapy (Figure S9a-10; Table S5-6). We observed significant downregulation of gene signatures related to neutrophils, skin-specific DCs, Langerhans cells, monocytes, IFN signaling, IL-21 and IL-12 pathways, complement system, inflammasome activation, apoptosis, the immunoproteasome and dermal Th2 cells—highlighting its broad immunosuppressive and anti-inflammatory effects. These effects were largely unique to dupilumab when compared to the effects of treatment with the other therapies tested in this meta-analysis. However, notably, the skin adhesion gene signature was consistently upregulated with all therapeutic agents, whereas the skin barrier signature remained unchanged with treatment (Figure S9-11; Table S5-6). In an additional study (GSE157194) comparing dupilumab treatment with the calcineurin inhibitor, cyclosporine, we found that dupilumab treatment led to greater transcriptomic improvement from baseline to month 3 (Figure S12). Both treatments reduced expression of cytokine-related signatures as well as monocyte and myeloid cell–associated pathways. However, dupilumab demonstrated a broader and more pronounced impact, showing greater suppression of Th1, Th2, Th17, IFN-related and anti-inflammatory gene signatures. In contrast, cyclosporine exerted a more modest and selective effect, downregulating monocytes/myeloid cell, IL12 complex, IL21 complex and inflammasome gene signatures.

The anti-IL22 antibody, fezakinumab, also induced modest, but detectable molecular shifts in LES of AD patients by week 12 of treatment (Figure S9b-10; Table S5-6). Endothelial and fibroblast signatures were upregulated, consistent with tissue remodeling, whereas the IL-17 complex was downregulated. Analysis of skin-specific T cell subsets revealed downregulation of both Th2 cell and Treg signatures with fezakinumab treatment (Figure S9b-10, Table S5-6).

The PDE4 inhibitor crisaborole exhibited a distinct molecular profile, perhaps related to its topical method of application (Figure S9c-10; Table S5-6). The TGF-β-associated fibroblast and unfolded protein gene signatures were decreased exclusively in crisaborole-treated samples. In addition, metabolic pathways, including the pentose phosphate pathway, TCA cycle and OXPHOS, were uniquely downregulated with crisaborole, pointing to broader immunometabolic reprogramming. As a second topical therapeutic agent, we examined the impact of the JAK1/Jak3/Tyk2/Syk inhibitor cedulatinib on gene signatures in AD patients (Figure S9d-10; Table S5-6). Like crisaborole, cedulatinib treatment decreased TNF signaling, anti-inflammation, cell cycle, and keratinocyte signatures, but had a broader impact beyond this, likely related to targeting numerous kinases. Specifically, cedulatinib also targeted many of the same cells/pathways as dupilumab, including neutrophils, monocyte/myeloid cells, IFN signaling, and complement genes.

Therapeutic analysis of the oral PDE4 inhibitor, apremilast, showed dose-dependent treatment effects in AD patients after 12 weeks of treatment (Figure S9e, 11). At the lower dose (30mg) of apremilast, the erythrocyte signature was uniquely decreased and the dermal CD8 T cell signature was increased as compared to the higher dose. At the 40 mg dosage, we observed downregulation of the keratinocyte, IL-17/IL-23 pathway, TNF signaling, apoptosis, cell cycle, glycolysis and proteasome gene signatures. The inflammasome and skin adhesion pathways were downregulated by both the 30 mg and 40 mg doses, indicative of common transcriptomic changes across dosages. A summary comparing and contrasting the various effects of each therapeutic agent on molecular profiles of AD patients can be found in Table S6.

### Dupilumab modulates blood immune profiles in AD patients based on molecular endotypes

To assess the systemic impact of dupilumab treatment on gene expression profiles, we analyzed GSVA scores of 32 immune and inflammatory gene signatures (Hubbard et al. 2023) in blood samples from 47 adult AD patients (Möbus et al. 2022), collected at baseline and 3 months after treatment (Figure 5a). Overall, treatment led to a reduction in the activity of multiple immune-related modules, including those associated with IFN, granulocytes, LDGs, monocytes, neutrophils, inflammasome, B cells, gd T cells and NK cells (Figure 5b).

**Figure 5:**
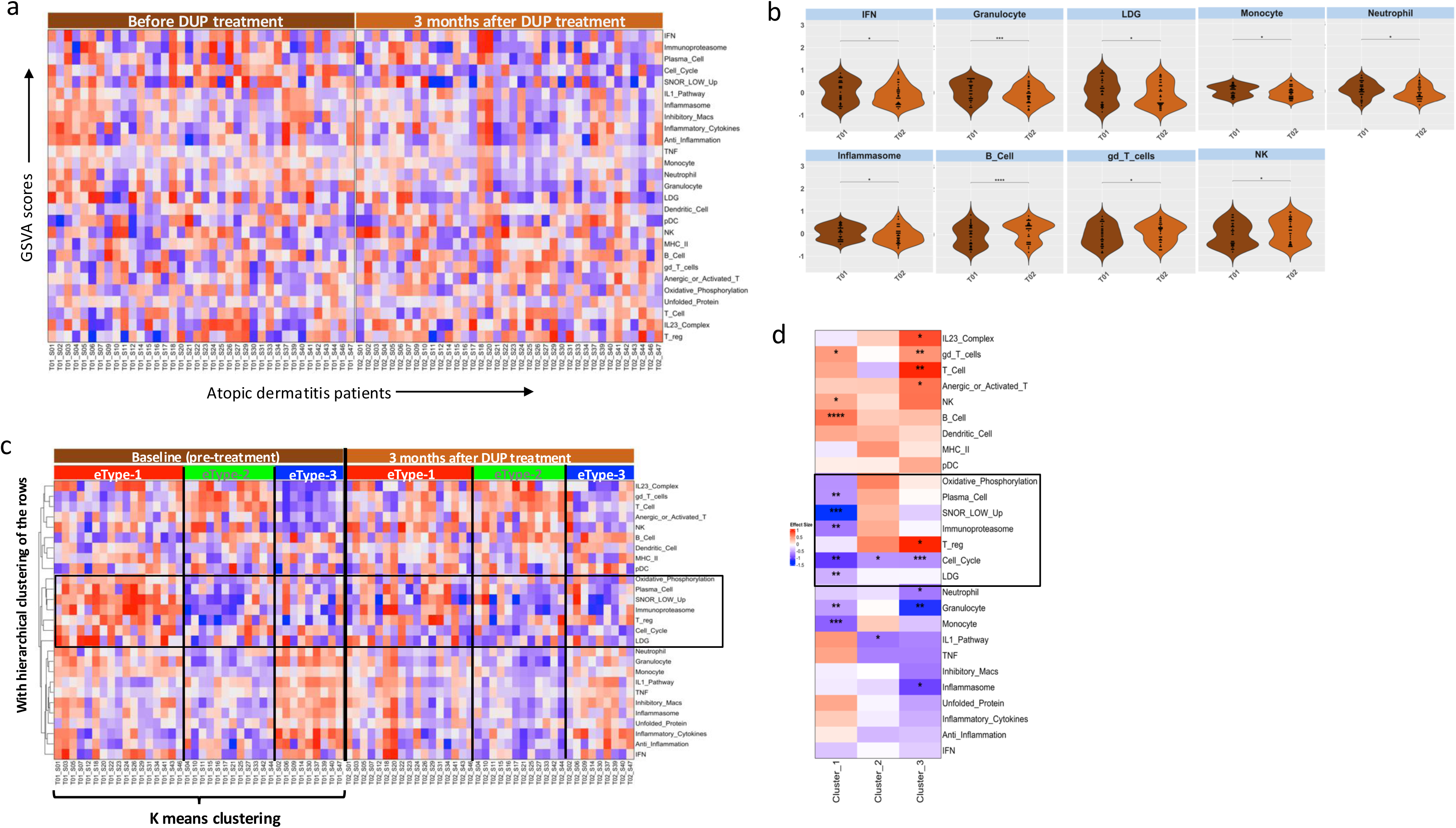
Transcriptomic changes in blood immune signatures of adult AD patients treated with dupilumab. **a**. Heatmap showing GSVA scores of 32 gene modules in blood from 38 patients at baseline and after 3 months of treatment. **b.** Violin plots illustrating significant changes in selected immune signatures before and after treatment (paired t-test). **c**. Heatmap from (a) reordered by k-means clustering of baseline GSVA scores and hierarchical clustering of rows, revealing three transcriptomic endotypes (eType-1, eType-2, eType-3). The same order is retained post-treatment to visualize endotype-specific shifts. **d.** Heatmap of Hedges’ g effect sizes showing the magnitude and direction of GSVA score changes within each endotype. *P < 0.05; **P < 0.01; ***P < 0.001; ****P < 0.0001.

To investigate inter-patient heterogeneity in treatment response, we carried out k-means clustering on baseline GSVA profiles and identified three transcriptomic endotypes (eType-1, eType-2, and eType-3), each exhibiting distinct immune activation patterns (Figure 5c). Notably, the extent of transcriptomic change after treatment varied by endotype. Hedge’s g effect size analysis revealed that eType-1 patients exhibited the most pronounced transcriptomic shifts, particularly in IFN, IL1, inflammasome, neutrophil, and monocyte signatures, whereas eType-2 and eType-3 showed more modest or selective changes (Figure 5d). These findings suggest that baseline immune endotypes influence the magnitude of molecular response to dupilumab and may offer a path toward stratifying patients for treatment optimization.

### The magnitude of transcriptomic normalization in LES and NLS skin following dupilumab treatment

To determine the extent to which dupilumab normalizes AD-associated transcriptomic abnormalities, we compared GSVA scores across LES, NLS and CTL skin at week 16 in both placebo and dupilumab treated AD patients (Figure 6, Figure S13). The gene signatures with a significant change in the magnitude of GSVA scores between groups are displayed as an effect size heatmap (Figure 6). In placebo-treated patients, LES skin remained enriched for immune activation signatures, including T cell subsets, IFN responses, IL-1 cytokines, and antigen presentation modules, whereas NLS skin exhibited downregulation of multiple signatures including keratinocytes, TGFB fibroblast, immunoproteasome, and dermal Th2 cells as compared to control skin. In contrast, LES skin of dupilumab-treated patients demonstrated broad suppression of immune and cytokine modules and restoration of skin barrier modules, although gene expression profiles were not restored to the level of CTL skin. Specifically, inflammatory signatures involving T cells, myeloid cells, and key cytokine pathways were diminished, whereas certain gene signatures, including Th17 cells, B cells, unfolded protein, IL1 cytokines and IL12 remained elevated in treated LES skin. NLS skin of dupilumab-treated patients also exhibited downregulation of inflammatory gene signatures, although to a lesser extent than observed in LES skin as there was also less elevation of these signatures at baseline. In addition, gene signatures of Th17 cells remained elevated, whereas skin barrier signatures were further decreased in dupilumab-treated NLS as compared to CTL skin.

**Figure 6:**
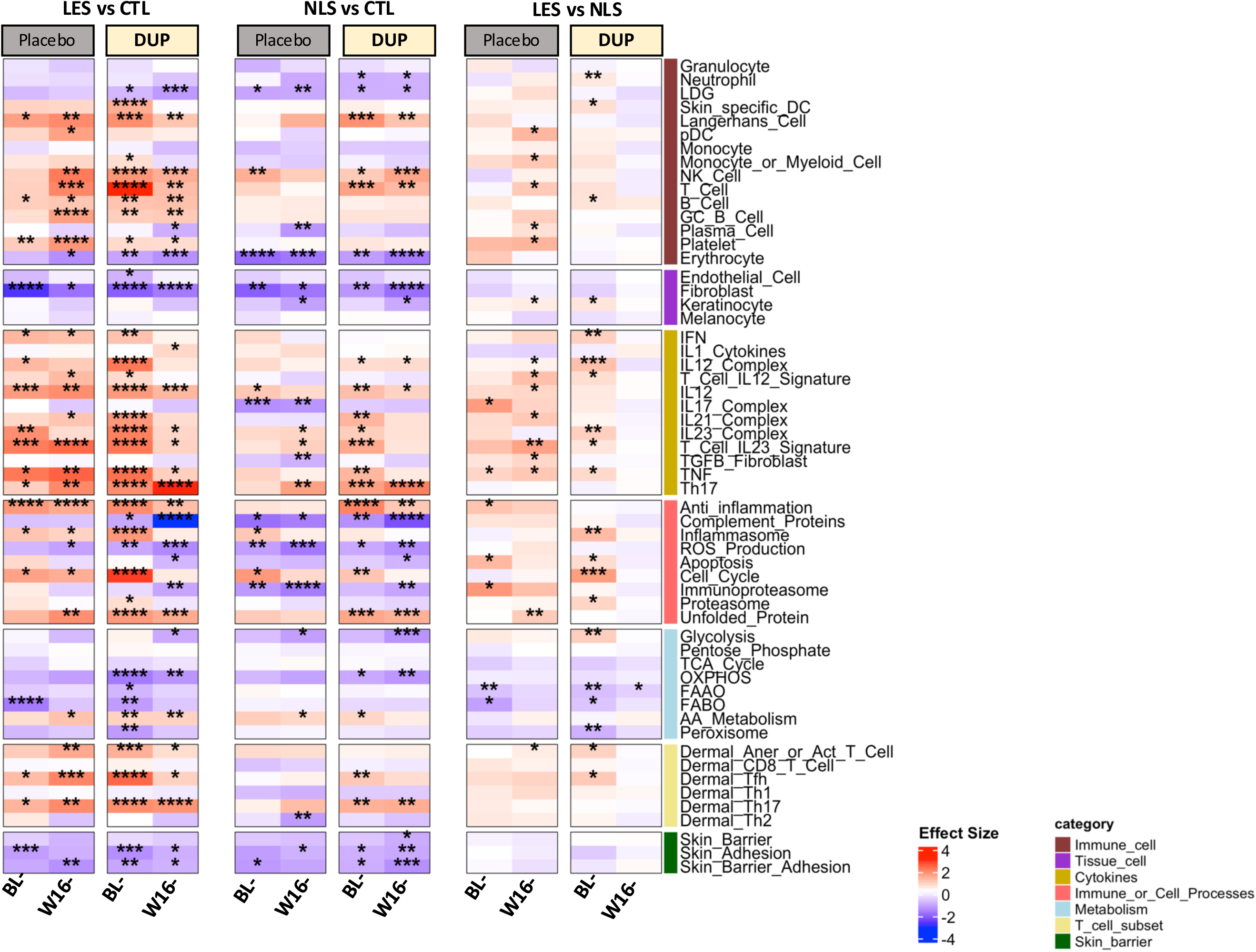
Statistical differences in molecular profiles of placebo-or dupilumab-treated LES and NLS skin compared with CTL. Heatmap showing Hedges’ g effect sizes representing the magnitude and direction of GSVA score changes across comparisons: LES vs CTL, NLS vs CTL, and LES vs NLS at baseline and Week 16 for both placebo and dupilumab groups. Red indicates higher, and blue lower enrichment relative to the comparator group. Asterisks denote significance based on Welch’s or paired t-tests (*P < 0.05; **P < 0.01; ***P < 0.001; ****P < 0.0001).

In order to assess the magnitude of change in GSVA scores over time in greater detail, we calculated Hedges’ g effect sizes for the contrasts LES vs CTL, NLS vs CTL, and LES vs NLS at baseline and week 16 of dupilumab treatment, and computed the delta (Δ Hedges’ g = Week 16 – Baseline) (Figure S14a-c). In the LES vs CTL comparison (Figure S14a), dupilumab-treated patients exhibited marked decreases in effect sizes across multiple disease-associated cell and pathway gene signatures as compared to placebo, in particular the T cell, IFN, IL12 complex, complement protein and cell cycle modules. When comparing NLS and CTL skin (Figure S14b), dupilumab induced modest but consistent reductions across inflammatory gene modules although the magnitude of change relative to placebo was reduced as compared to LES skin. The LES vs NLS comparison (Figure S14c) verified the increased magnitude of anti-inflammatory changes induced by dupilumab treatment in LES relative to NLS skin. Interestingly, we also observed increased enrichment of the metabolism and skin adhesion signatures specifically in LES but not NLS skin with dupilumab treatment. Dupilumab treatment induced broad transcriptomic normalization in LES skin, with inflammatory and cytokine pathways markedly suppressed and profiles approaching but not reaching those of CTL. In NLS skin, dupilumab produced more modest but consistent reductions in residual immune activation, indicating that while both compartments benefit, the magnitude of change is most pronounced in affected, LES skin. Together, these findings highlight dupilumab’s capacity to mitigate AD-associated abnormalities across both LES and NLS skin, with the greatest impact in diseased tissue.

## DISCUSSION

To deconvolute the heterogeneity of AD and identify common molecular features, we carried out a comprehensive transcriptomic analysis of LES and NLS skin from adult AD patients, benchmarked against CTLs, and further compared adult with pediatric AD, PSO and NME. We complied a consensus molecular profile of adult AD that was consistent across the majority of the gene expression datasets analyzed in this study. Notable changes in LES skin included enrichment of gene signatures of B and T cells and the cytokines IL-12, IL-23, and TNF as well as de-enrichment of skin barrier-associated gene signatures. This result is supported by molecular and clinical studies in humans and mouse models, which indicate that AD is primarily, but not exclusively, driven by a Th2 response, and may also involve contributions from Th1, Th22, and Th17 immune cell and cytokine responses leading to or as a result of increased skin barrier permeability (Facheris et al. 2023; Kim et al. 2019). In summary, although we found evidence for multiple types of immune responses in individual AD datasets, only IL-12, IL-21, and IL-23 cytokine-related gene signatures were consistently elevated in LES skin across all datasets suggesting that these are common drivers of AD pathogenesis that could be targeted for therapeutic intervention in the majority of AD patients.

We observed similar changes in inflammatory and skin barrier gene signatures in adult NLS AD skin, although the magnitude was diminished and the heterogeneity across datasets was increased as compared to LES skin. This is consistent with previous work in which NLS skin was found to be genomically and histologically distinct from normal skin, but with greater variability in immunologic involvement than LES skin (Suárez-Fariñas et al. 2011). In summary, investigation of NLS skin has the potential to provide information about the early development of AD lesions, but requires additional analysis of paired LES and NLS skin.

Our analysis also revealed clear differences between adult and pediatric AD. Lesional skin of pediatric AD was characterized by more restricted immune dysregulation, with IL-1–driven inflammation emerging as a key feature, significant enrichment of Th2 and Th17 cells, and an absence of significant barrier-related defects. This result aligns with prior pediatric studies of cytokine responses in cord blood showing strong type 2 and variable Th17 polarization, but limited Th1 activation as risk factors for development of AD (Brunner et al. 2019a; Herberth et al. 2010; Van Der Velden et al. 2001) In addition, several comparative studies of pediatric and adult AD skin biopsies implementing immunohistochemical and transcriptomic analyses have described a shared Th2 program across ages but noted adult-specific amplification of Th1/IFN-γ responses and more pronounced barrier defects in adult AD patients (Brunner et al. 2018a; Renert-Yuval et al. 2021). Direct comparison of healthy adult and pediatric skin biopsies revealed similar levels of heterogeneity in enrichment of most inflammatory gene signatures, although some inherent differences in other gene signatures emerged. Notably, dermal T cell signatures were generally enriched, whereas skin barrier and adhesion signatures were de-enriched in healthy pediatric as compared to healthy adult skin. Thus, age-dependent molecular profiles of healthy skin may, to some extent, inform the nature of disease pathology in individuals with AD, although it is important to note that all gene signature enrichments observed in LES AD skin were determined relative to their corresponding healthy controls.

Early studies in which AD was induced through experimental allergic sensitization proposed that AD pathology was driven by a biphasic immune response with an acute phase driven by a type-2/Th2 response and a chronic phase sustained by continued type-1/Th1 cytokine-mediated inflammation, both of which contribute to impaired skin barrier functionality (Grewe et al. 1995; Thepen et al. 1996). In the context of this system, the involvement of Th2 responses in pediatric and Th1 responses in conjunction with decreased skin barrier gene expression in adult AD suggests that pediatric AD represents early/acute phase disease whereas adult AD is representative of acute as well as late/chronic phase AD. In addition, the relative de-enrichment of skin barrier and adhesion gene signatures in healthy pediatric as compared to healthy adult skin indicates that this barrier is not as developed in children as compared to adults. These findings suggest that the immunopathology of AD evolves with age, with pediatric disease driven by acute, Th2-dominant responses, which have not yet resulted in skin barrier defects, whereas adult disease reflects a convergence of Th2/Th1/Th17 responses and more prominent barrier dysfunction.

Cross-disease comparisons underscore shared inflammation, but distinct dominant drivers. All three diseases showed immune and cytokine pathway enrichment in LES skin versus NLS skin, but the magnitude and composition of these changes varied. AD was marked by strong immune dysregulation coupled with suppression of skin barrier signatures. These abnormalities were more pronounced than in PSO, underscoring a broader disruption of immune–metabolic balance in AD. In line with previous findings, PSO was characterized by increased enrichment of keratinocytes and metabolic activation (Pasquali et al. 2019, Martinez et al. 2022). NME exhibited unique immune features and demonstrated the strongest degree of overall immune activation both in the number of enriched immune cell gene signatures as well as the increased degree of enrichment. In summary while immune activation is a common thread across these diseases, the dominant drivers differ—immune dysregulation and barrier impairment in AD, metabolic rewiring in PSO (Sarandi et al. 2023), and broad, heightened immune responses in NME including greater enrichment of pro-inflammatory cells and cytokines.

Longitudinal analysis of untreated AD skin further highlighted dynamic variation in immune pathways over time. While control skin was largely stable, LES skin displayed fluctuating activation of multiple inflammatory pathways at different stages of follow-up, suggesting that immune drivers of disease may shift over time. NLS skin also showed temporal changes although to a lesser extent than LES skin, underscoring the systemic nature of disease-associated abnormalities. Notably, these molecular changes did not correlate with clinical severity scores, emphasizing that transcriptomic measures capture dimensions of disease activity or standard of care therapy not reflected in clinical scoring systems. To address this gap, we developed ECZECIS, a composite transcriptomic score that integrates common disease-associated molecular features across AD datasets into a system in which the features contributing most to each patient’s disease can be monitored in real time. Notably, although ECZECIS did not correlate with clinical severity in untreated patients, it strongly aligned with both EASI and SCORAD during dupilumab treatment, highlighting its utility as a molecular biomarker of therapeutic response.

Of therapeutic interventions, dupilumab, targeting the Th2 cytokines IL4/IL13, exerted the most extensive transcriptomic alterations, broadly suppressing Th1, Th2, Th17, IFN, myeloid, and apoptotic pathways, and aligning LS skin profiles more closely with NLS skin (Hamilton et al. 2021). These findings are consistent with prior clinical studies showing that dupilumab progressively shifts molecular phenotype from LES to NLS (Guttman-Yassky et al. 2019) and suppresses mRNA expression of genes of type 2 inflammation. Comparisons with the broad immunosuppressive agent cyclosporine revealed that while both agents reduced cytokine and myeloid cell signatures, dupilumab achieved a broader spectrum of immune suppression, consistent with its clinical efficacy (Möbus et al. 2021). Other agents, including fezakinumab (anti-IL22), crisaborole (topical PDE4 inhibitor), cedulatinib (topical Jak kinase family inhibitor) and apremilast (oral PDE4 inhibitor), induced more selective transcriptional shifts, reflecting their narrower mechanisms of action as has also been noted in clinical and molecular studies of these drugs (Brunner et al., 2017; Czarnowicki et al., 2018; Piscitelli et al., 2021). Importantly, shared transcriptional targets across these drugs—such as IL-23, TNF, and proteasome pathways—suggest common molecular nodes of therapeutic benefit, despite differences in molecular targets of each agent (Bissonnette et al. 2019; Brunner et al. 2019b; Samrao et al. 2012; Schafer et al. 2019).

Across all comparisons in our study, barrier-related pathways consistently emerged as a key differentiator between cohorts. To evaluate different aspects of skin barrier function we assessed expression of transcripts involved in skin adhesion, or the membranous junctions between skin cells, separate from the canonical barrier transcripts, most notably filaggrin (FLG). Adult AD patients showed uniform downregulation of skin barrier/adhesion signatures in both LS and NL skin, whereas pediatric AD patients lacked significant barrier abnormalities but did exhibit decreases in adhesion transcripts, indicating that pediatric AD may represent early stages of skin barrier dysfunction that is a prominent feature of adult disease. This is also consistent with previous findings that pediatric AD cohorts exhibited early lipid and metabolic abnormalities rather than global suppression of barrier transcripts including FLG (Brunner et al. 2018b; Esaki et al. 2016).

When comparing barrier involvement in AD to other inflammatory dermatoses, PSO was notable for an upregulation of barrier-associated modules along with downregulation of the skin adhesion signature, whereas both AD and NME exhibited marked downregulation of both skin barrier and adhesion genes. These findings align with earlier literature establishing barrier impairment as a hallmark of AD pathophysiology, driven by both structural protein deficiencies and immune-mediated disruption (Cork et al. 2009; Elias 2008; Weidinger and Novak 2016). In addition, as null mutations in FLG remain a significant risk factor for the development of AD, the specific involvement of the skin barrier signature in AD but not PSO highlights the pivotal role of skin barrier integrity in AD (Bin and Leung 2016). Therapeutic analysis further revealed that the skin barrier module remained largely unchanged across treatments, highlighting the persistence of barrier impairment even with effective immunomodulation.

Finally, blood-based profiling (Möbus et al. 2022) revealed that dupilumab reduces systemic immune activation across multiple cell types and cytokine pathways, and that baseline molecular endotypes influence the magnitude of response. Patients with elevated baseline IFN, inflammasome and myeloid activation (eType-1) experienced the most pronounced transcriptomic improvements, supporting the concept of molecular endotyping as a strategy for treatment stratification (Hubbard et al. 2023a).

In summary, AD is characterized by widespread immune skewing and barrier defects that differ by age, diverge across related skin diseases, fluctuate over time, and are variably shaped by therapy—most broadly by dupilumab. Incorporating transcriptomic scoring such as ECZECIS with clinical assessment captures treatment-related improvement that routine scores may miss and, together with blood-based endotypes, points toward patient stratification for targeted interventions. Key limitations include reliance on a single pediatric and a single longitudinal dataset and heterogeneous study designs, underscoring the need for validation in larger, age-stratified, prospective cohorts. Moving forward, embedding standardized transcriptomic endpoints in clinical studies and testing endotype-guided treatment strategies may enable more precise monitoring and truly personalized care for patients with AD.

## MATERIALS AND METHODS

### Datasets

This study utilized ten publicly available microarray (GSE130588 Guttmann-Yassky et al. 2019, GSE32924 Suarez-Farinas et al. 2011, GSE107361 Brunner et al. 2018, GSE99802 Brunner et al. 2019, GSE133477 Bissonnette et al. 2019, GSE120899 Simpson et al. 2019) and bulk RNA-seq datasets (GSE121212 Tsoi et al. 2020, GSE157194 Möbus et al. 2021, GSE193309 Hu et al. 2023, GSE208405 Möbus et al. 2022, GSE141570 Piscitelli et al. 2021) retrieved from the Gene Expression Omnibus (Table S1).

### Data analysis and statistics

Processing and analysis of gene expression datasets and development of the ECZECIS transcriptional scoring system was carried out as previously described (Shrotri et al. 2024). **Detailed methods are available in Supplementary Materials.**

## DATA AVAILABILITY STATEMENT

All datasets are publicly available from the GEO database as detailed in Table S1.

## CONFLICT OF INTEREST STATEMENT

The authors declare no conflicts of interest.

## ACKNOWLEDGMENT

We thank our colleagues at AMPEL BioSolutions for their insight and contribution to scientific discussions related to this work. This work was supported by the RILITE Foundation.

## AUTHOR CONTRIBUTIONS

Conceptualization: S.S., A.D., and P.L. Methodology: S.S., A.D., and P.B. Validation: S.S. and A.D. Formal Analysis: S.S. Investigation: S.S. and A.D. Data curation: S.S. and P.B. Writing-original draft: S.S. and A.D. Writing-review and editing: S.S., A.D., and P.L. Visualization: S.S. and A.D. Supervision: A.G., P.B., and P.L. Funding acquisition: A.G. and P.L.

## TABLES

Supplementary Tables S1-S6

## Notes

### Competing Interest Statement

The authors have declared no competing interest.

