## Supplemental Materials for "Comprehensive Transcriptomic Analysis of Atopic Dermatitis Patients Documents the Spectrum of Molecular Abnormalities and the Response to Treatment"

### SUPPLEMENTARY MATERIALS

#### Data preprocessing

For microarray datasets, CEL files were obtained from GEO and normalized using the GeneChip Robust Multiarray Average (GCRMA) method, with normalization and sample processing performed independently for each dataset. For bulk RNA-seq datasets, FASTQ files were downloaded from GEO, subjected to quality control with FASTQC, aligned to the human reference genome (GRCh38) using STAR, and quantified with featureCounts. Count matrices were imported into DESeq2, low-abundance genes were filtered with HTSFilter, and size-factor normalization was applied. For downstream analyses, normalized expression values were obtained using the normTransform() function in DESeq2, producing log2-scaled normalized counts.

#### Gene set variation analysis

GSVA, a non-parametric, unsupervised method that computes sample-wise gene set enrichment scores (Hänzelmann et al. 2013), was performed using the gsva R package (version 1.40). Genes with interquartile range (IQR) > 0 were retained as input. For skin datasets, GSVA was conducted using 56 curated gene signatures (Martínez et al. 2022; Shrotri et al. 2024; Table S2) grouped into seven functional categories: immune cells, tissue cells, immune cell processes, cytokines, metabolism, T cell subsets, and skin barrier function. For blood datasets, 32 previously established immune signatures (Hubbard et al. 2023b) were used.

#### Statistical analysis

Statistical differences of GSVA scores between various groups were calculated using Welch's t test for unpaired samples, paired t test for paired samples or repeated measures of ANOVA for paired longitudinal samples. Bonferroni correction for p values was applied for repeated measures of

ANOVA. The magnitude of difference (effect size) was estimated using Hedge's g as calculated below:

$$g = \frac{\overline{x_1} - \overline{x_2}}{\sqrt{\frac{(n_1 - 1) * s_1^2 + (n_2 - 1) * s_2^2}{n_1 + n_2 - 2}}}$$

**where,**

$\overline{x_1}$ *and* $\overline{x_2}$ = *cohort* 1 *mean and cohort* 2 *mean respectively*

$n_1$ *and* $n_2$ = *cohort* 1 *size and cohort* 2 *size respectively*

$s_1^2$ *and* $s_2^2$ = *cohort* 1 *variance and cohort* 2 *varaince respectively*

cohort 1 and cohort 2 could be either LS sample at a particular timepoint, or LS / NL sample at baseline. All the statistical analysis was carried out in using effectSize (version 0.8.1) and stats (version 3.6.2) library in R. For visualization of GSVA scores and effect sizes ComplexHeatmap library and for the violin plots ggplot2 library in R was used.

**<u>Logistic Regression and ECZECIS (Eczema Cell and Immune Score)</u>**

Logistic regression with ridge penalty was carried out on GSVA scores of LS and CTL samples at baseline. Ridge penalization helped in coefficient shrinkage and in prevention of multicollinearity. Ridge penalized coefficients were then used to derive ECZECIS (ECZEma Cell and Immune Score) for each patient. Briefly, to calculate ECZECIS for each patient, GSVA scores were multiplied by coefficients and summed to generate a raw score. Raw scores were then normalized by adding a minimum score. ECZECIS scores approximately ranged from 0 to 15. ECZECIS scores were correlated to EASI or SCORAD using Pearson correlation, and statistical differences between EASI scores or SCORAD or ECZECIS scores across various timepoints were estimated using repeated measures of ANOVA with Bonferroni correction for p values.

**Machine Learning Classification of Lesional (LES) vs. Control Skin (CTL):**

GSVA enrichment scores of baseline LES AD skin samples and CTL samples were used as input features for machine-learning classification. Five supervised machine-learning algorithms were implemented using Python (scikit-learn v1.3.0): Logistic Regression (LR), k-Nearest Neighbors (KNN), Random Forest (RF), Naïve Bayes (NB), and Support Vector Machine (SVM). Model performance was evaluated using standard classification metrics, including sensitivity, specificity, precision, F1-score, Cohen's kappa coefficient, Matthews correlation coefficient (MCC), overall accuracy, and the area under the ROC curve (AUC). ROC curves were generated by plotting the true-positive rate against the false-positive rate across probability thresholds. To identify features contributing most strongly to LES vs. CTL discrimination, we extracted Gini importance scores from the trained Random Forest classifier. Feature importance values were ranked and visualized as a bar plot using ggplot2 package in R. All analyses were performed in Python 3.10 using scikit-learn, NumPy, pandas.

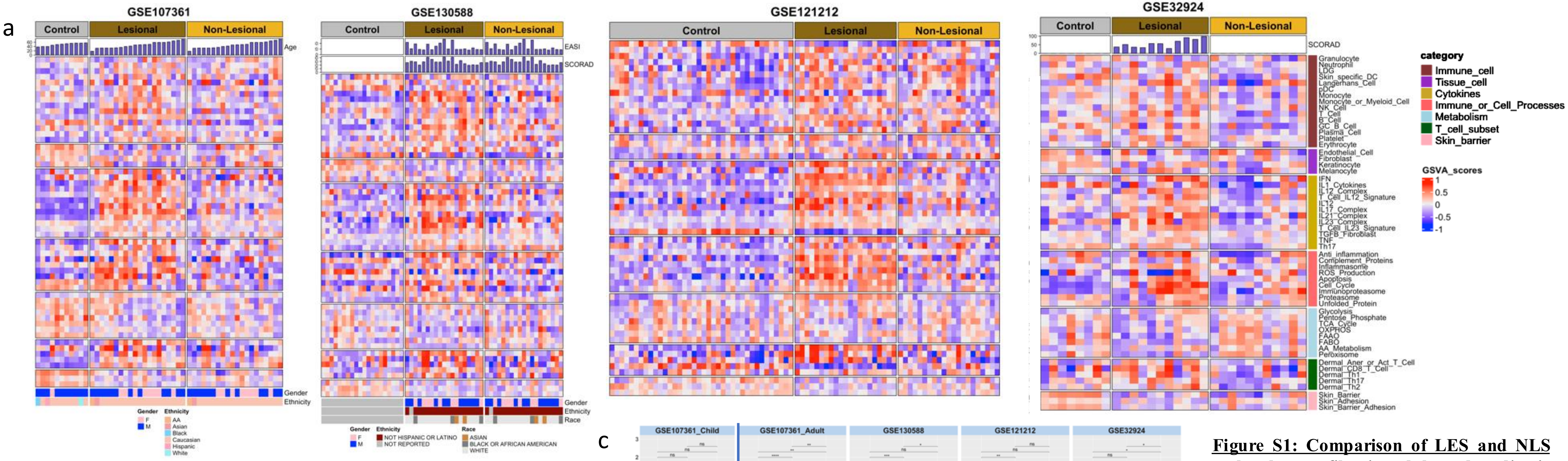


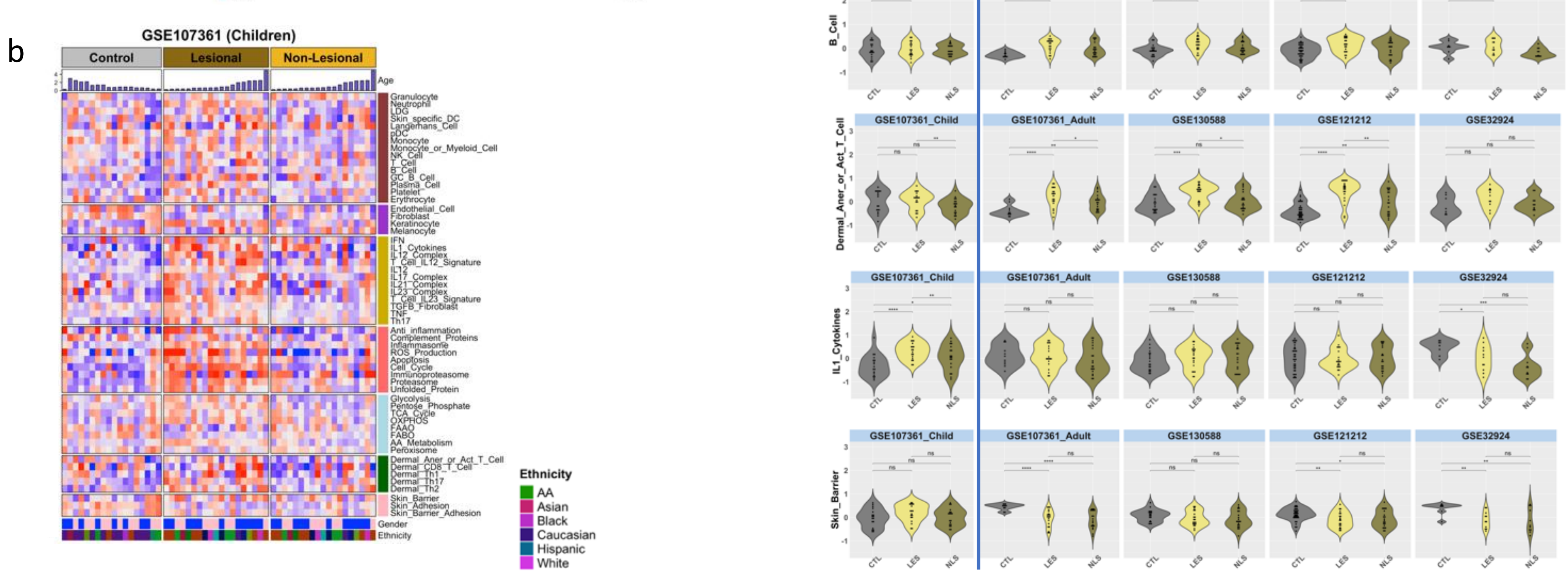


**Figure S1: Comparison of LES and NLS molecular profiles in adult and pediatric AD patients versus healthy controls at baseline across multiple datasets.** **a.** Heatmap showing GSVA scores for LES and NLS skin from adult AD patients and controls. **b.** Heatmap showing GSVA scores for pediatric AD and CTL skin. 56 gene signatures used for GSVA are grouped into seven categories: Immune cells, Tissue cells, Cytokines, Immune cell processes, Metabolism, T-cell subsets, and Skin barrier. **c.** Violin plots showing significant differences in B-cell, anergic/activated T-cell, IL1 cytokine, and skin barrier signatures between LES, NLS, and CTL skin. Dots represent individual GSVA scores; violin shapes indicate score distribution. *P < 0.05; **P < 0.01; ***P < 0.001; ****P < 0.0001.

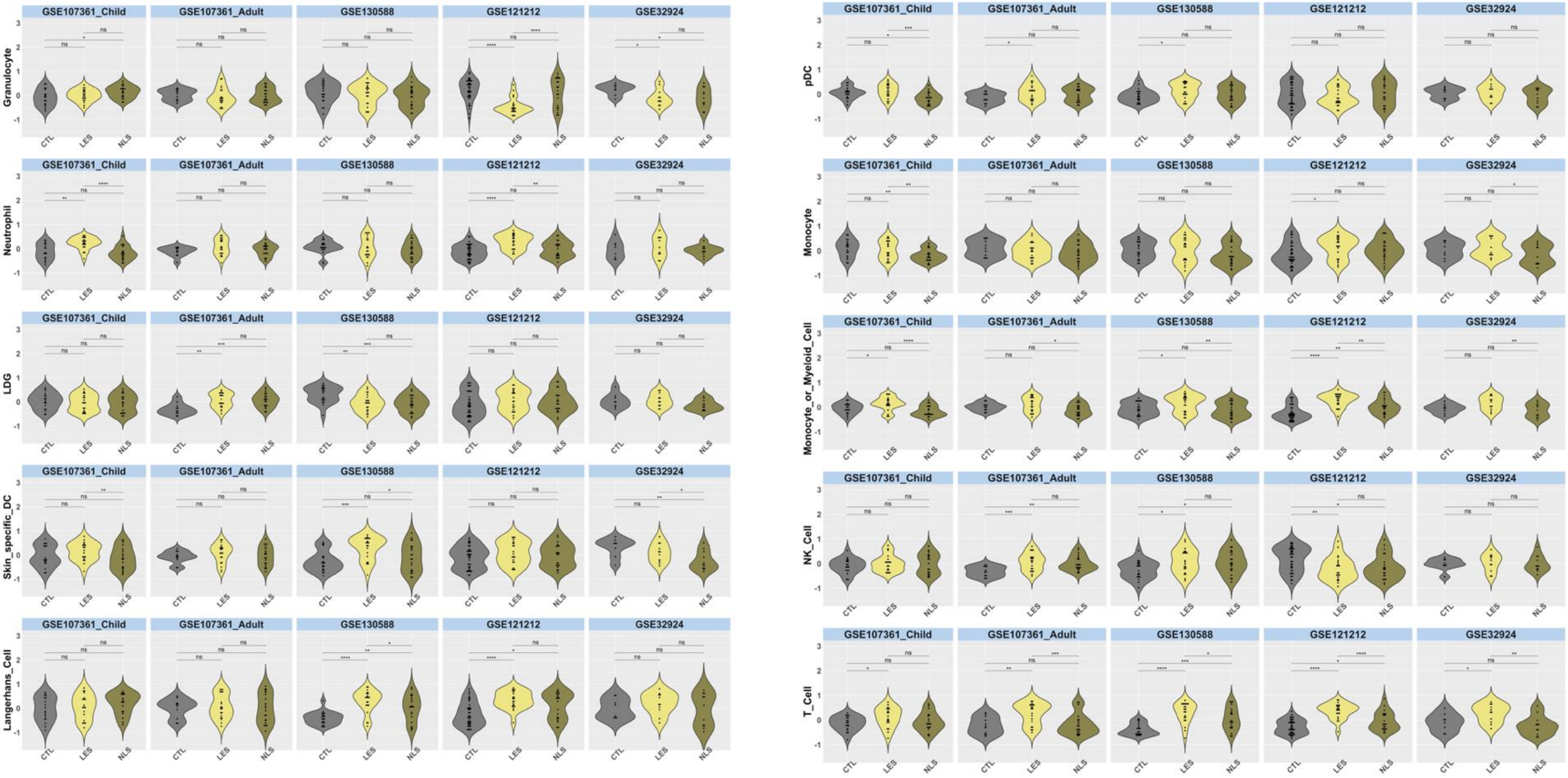


**Figure S2. Violin plots displaying changes in the GSVA scores in LES and NLS skin from AD patients compared to healthy controls across multiple datasets:** Violin plots display GSVA scores for 56 curated gene modules across five independent transcriptomic datasets: GSE107361_Child, GSE107361_Adult, GSE130588, GSE121212, and GSE32924. Each plot shows distribution of GSVA scores in CTL, LES, and NLS skin. Dots represent individual samples, and violin shapes indicate score distributions. Statistical comparisons were performed using paired t test, with significance annotated as follows: ns = not significant, *$p < 0.05$, **$p < 0.01$, ***$p < 0.001$, ****$p < 0.0001$.

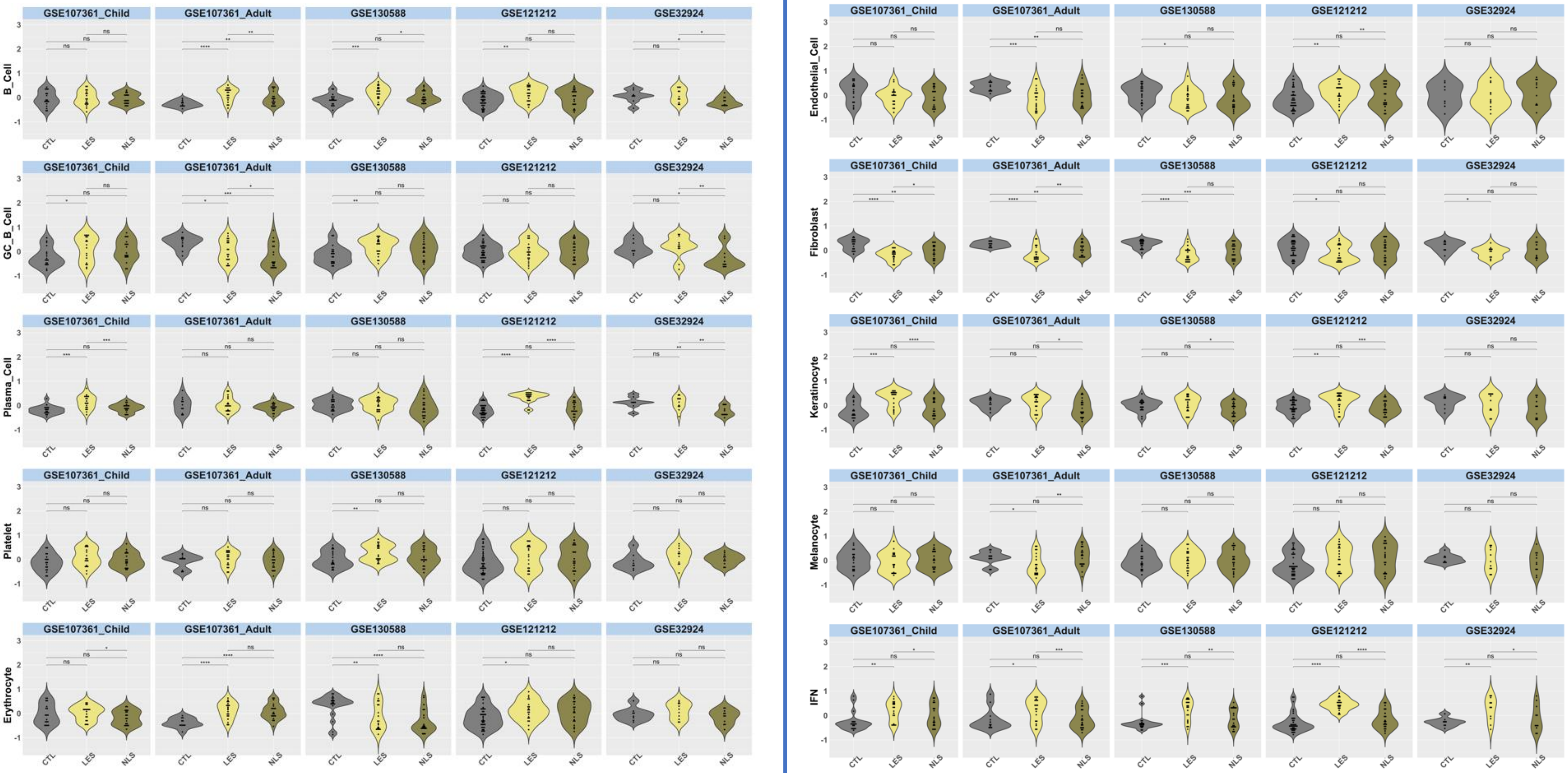
GSE107361_Child
GSE107361_Adult
GSE130588
GSE121212
GSE32924
B_Cell
GC_B_Cell
Plasma_Cell
Platelet
Erythrocyte
Endothelial_Cell
Fibroblast
Keratinocyte
Melanocyte
IFN
CTL
LES
NLS

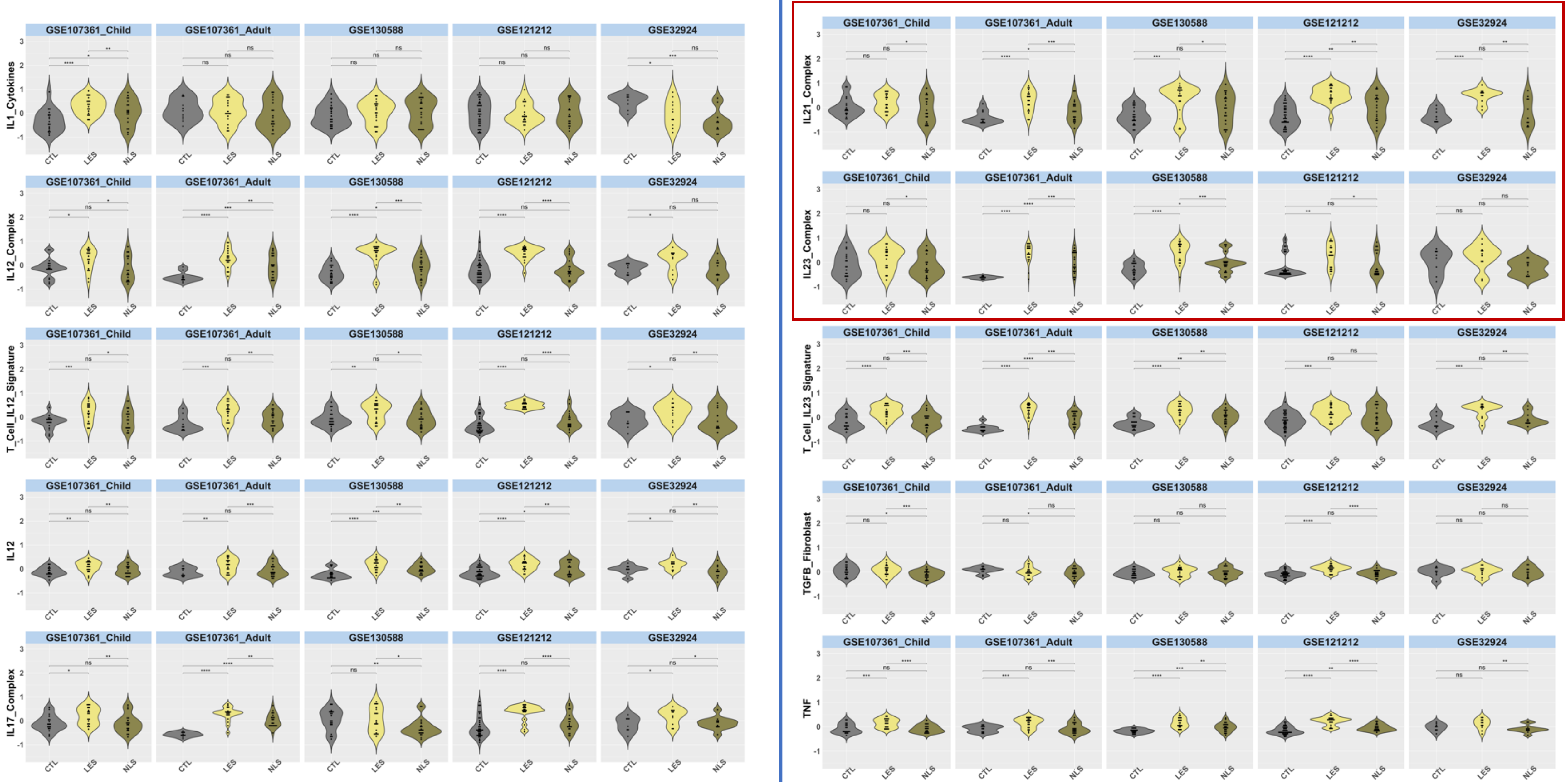

GSE107361_Child
GSE107361_Adult
GSE130588
GSE121212
GSE32924
IL1_Cytokines
IL12_Complex
T_Cell_IL12_Signature
IL12
IL17_Complex
IL21_Complex
IL23_Complex
T_Cell_IL23_Signature
TGFB_Fibroblast
TNF
CTL
LES
NLS

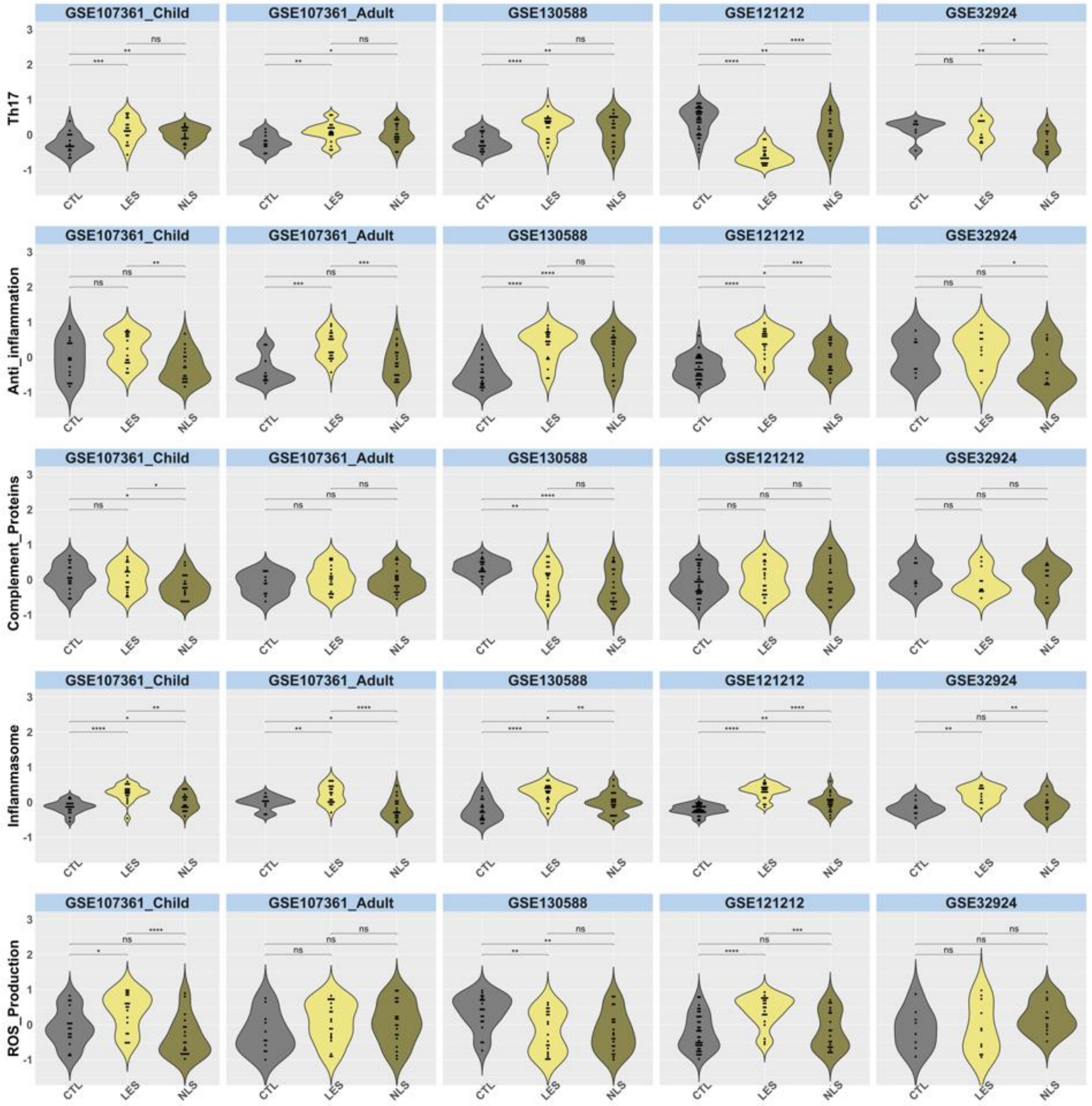
GSE107361_Child
GSE107361_Adult
GSE130588
GSE121212
GSE32924
Th17
Anti_inflammation
Complement_Proteins
Inflammasome
ROS_Production
CTL
LES
NLS

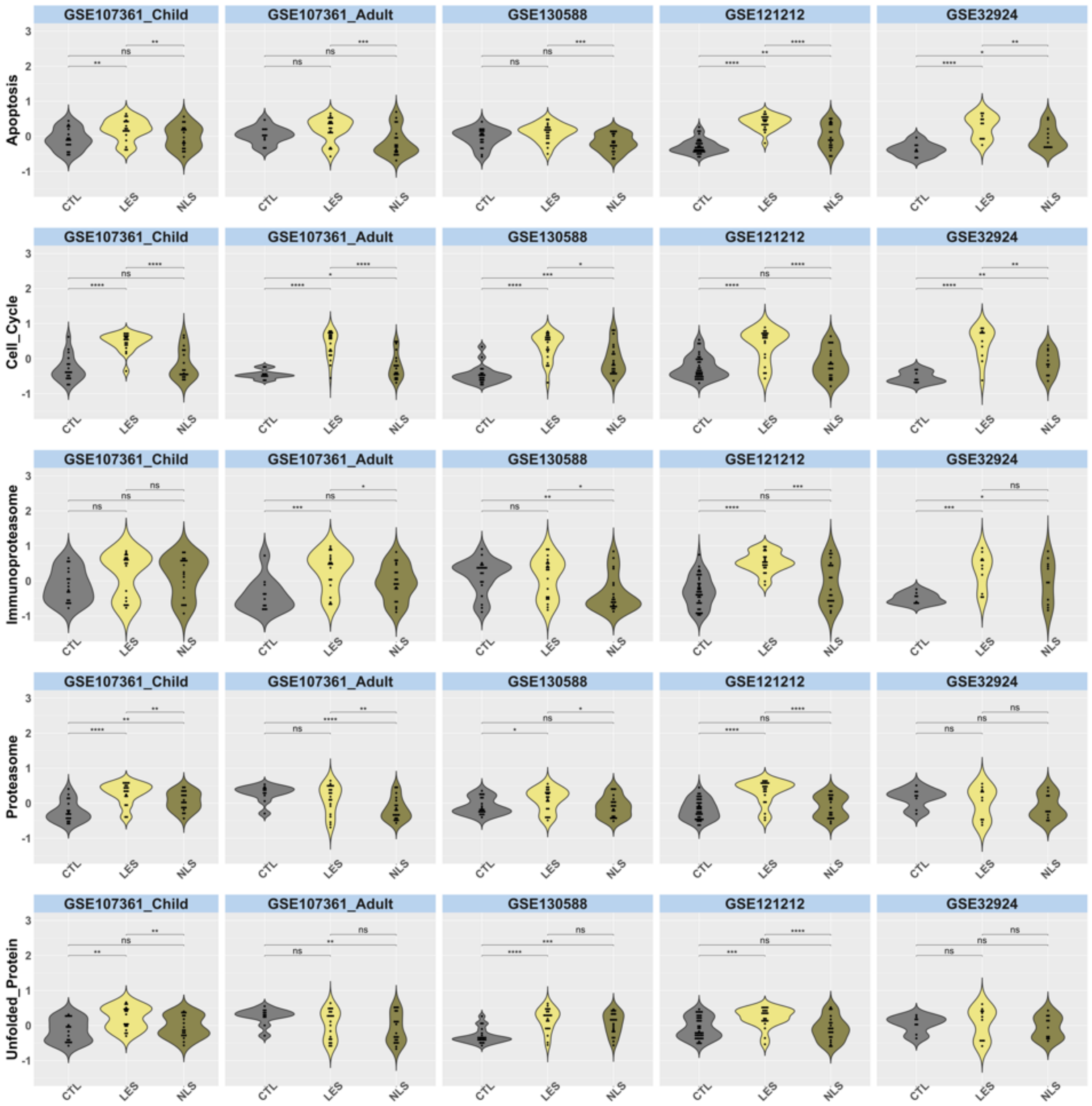
GSE107361_Child
GSE107361_Adult
GSE130588
GSE121212
GSE32924
Apoptosis
Cell_Cycle
Immunoproteasome
Proteasome
Unfolded_Protein
CTL
LES
NLS

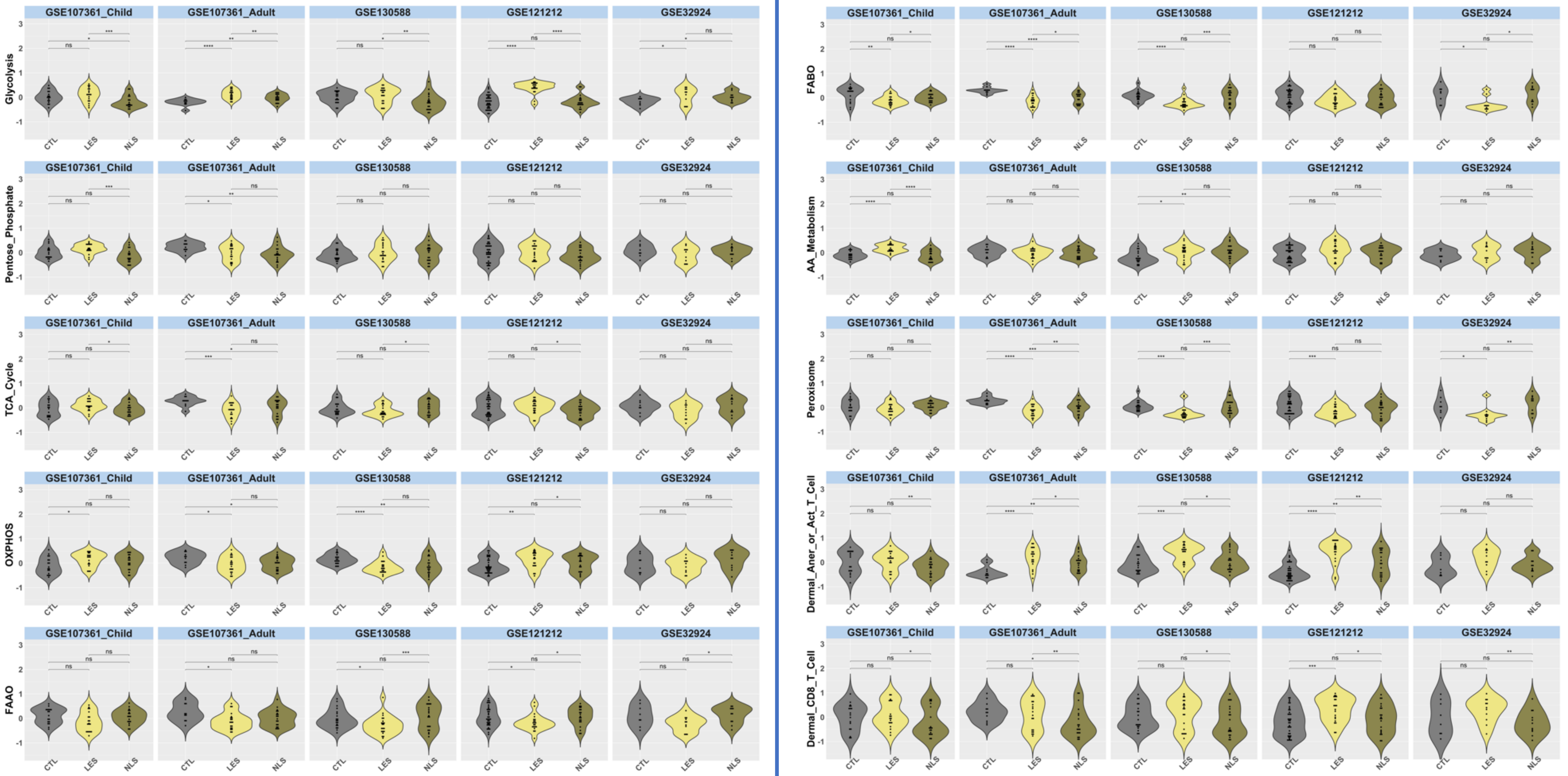
GSE107361_Child
GSE107361_Adult
GSE130588
GSE121212
GSE32924
Glycolysis
Pentose_Phosphate
TCA_Cycle
OXPHOS
FAAO
FABO
AA_Metabolism
Peroxisome
Dermal_Aner_or_Act_T_Cell
Dermal_CD8_T_Cell
CTL
LES
NLS

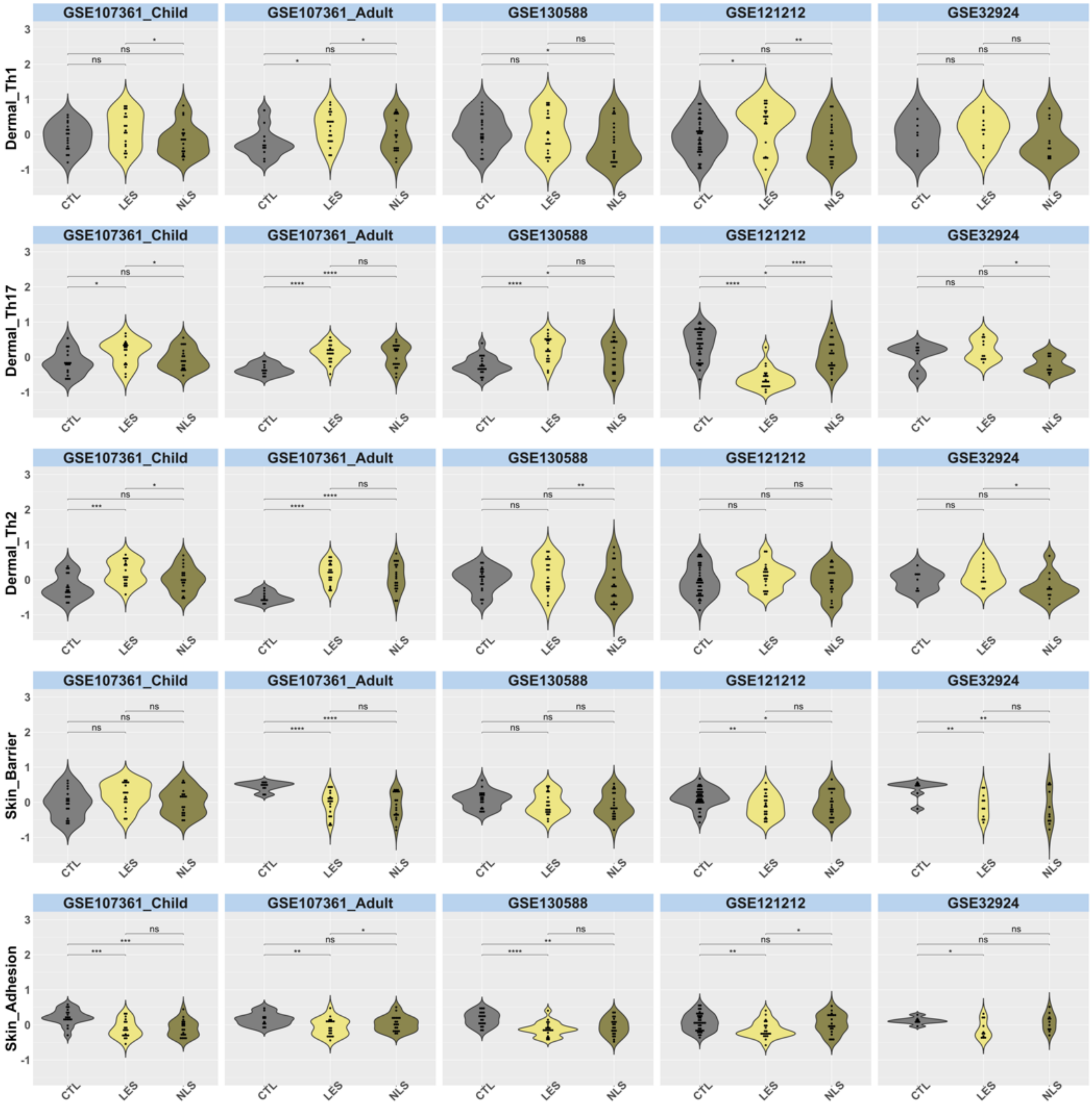

GSE107361_Child
GSE107361_Adult
GSE130588
GSE121212
GSE32924
Dermal_Th1
Dermal_Th17
Dermal_Th2
Skin_Barrier
Skin_Adhesion
CTL
LES
NLS

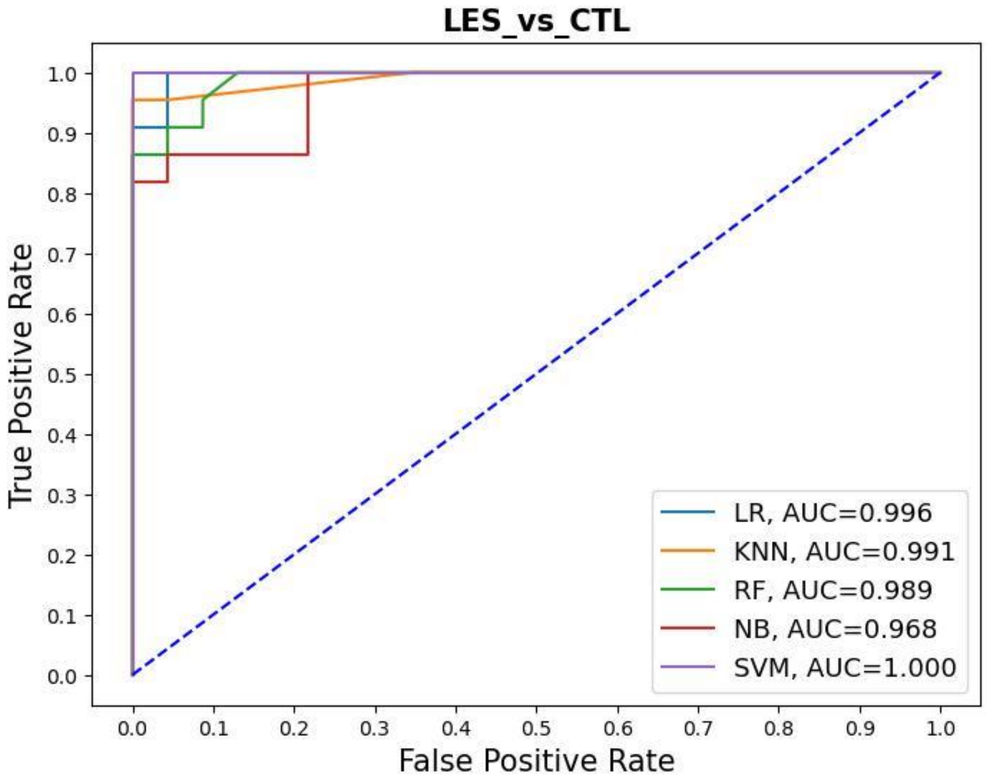


| Classifier | Sensitivity | Specificity | Cohen_Kappa_Score | Matthew Coeff | Precision | f-1 score | Accuracy |
|---|---|---|---|---|---|---|---|
| LR | 0.91 | 1 | 0.91 | 0.91 | 1 | 0.95 | 0.96 |
| KNN | 0.82 | 1 | 0.82 | 0.83 | 1 | 0.90 | 0.91 |
| RF | 0.86 | 1 | 0.86 | 0.87 | 1 | 0.92 | 0.93 |
| NB | 0.82 | 0.96 | 0.77 | 0.78 | 0.95 | 0.88 | 0.89 |
| SVM | 0.86 | 1 | 0.86 | 0.87 | 1 | 0.92 | 0.93 |

**Figure S3. Performance of multiple machine learning classifiers distinguishing LES from CTL skin a.** Receiver operating characteristic (ROC) curves for five machine-learning classifiers—Logistic Regression (LR), k-Nearest Neighbors (KNN), Random Forest (RF), Naïve Bayes (NB), and Support Vector Machine (SVM)—trained to distinguish LES from CTL skin using GSVA signature scores. **b.** Summary table of classifier performance metrics.

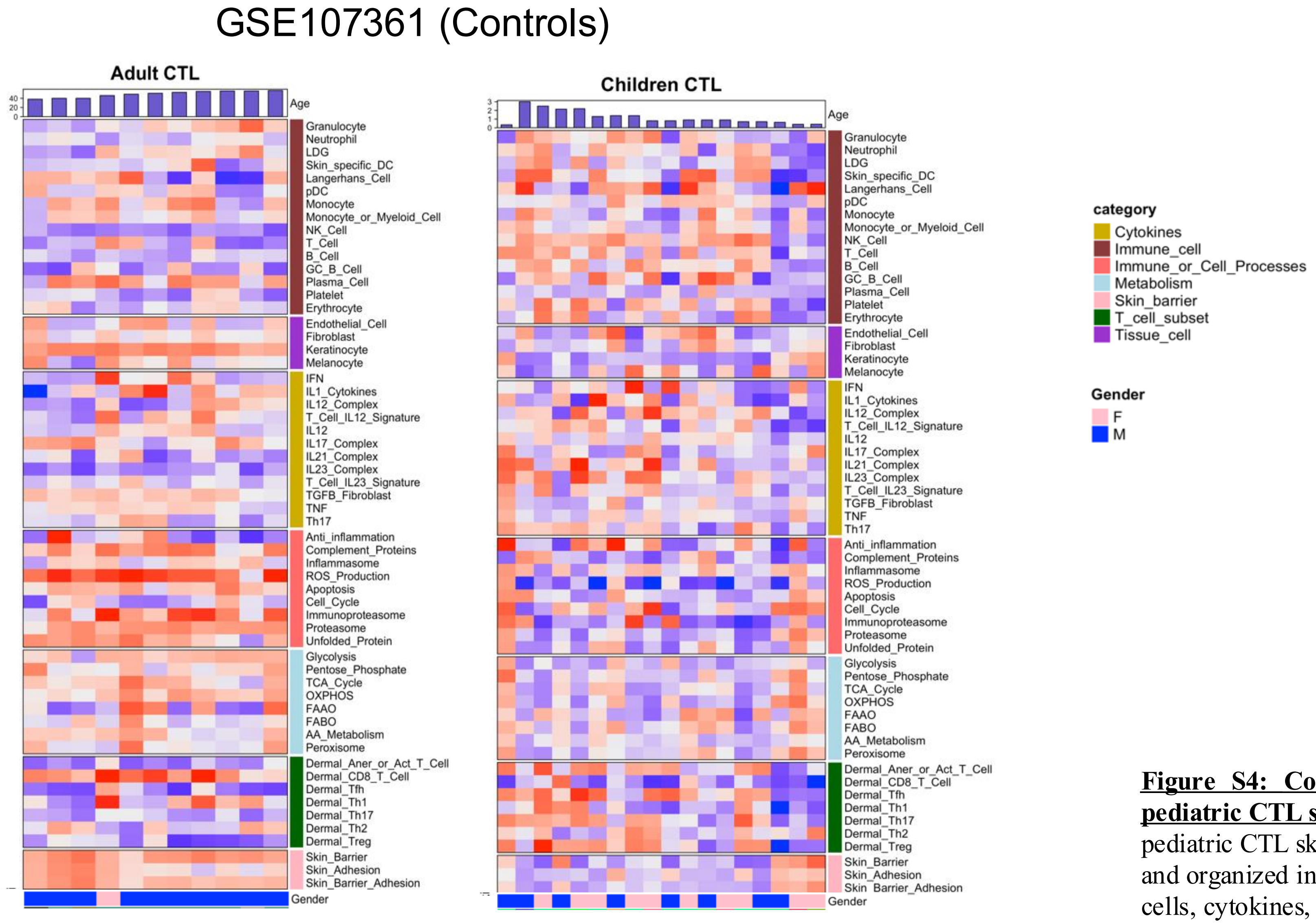


**<u>Figure S4: Comparison of molecular profiles in adult and pediatric CTL skin:</u>** Heatmap depicting GSVA scores for adult and pediatric CTL skin samples. A total of 58 gene signatures were used and organized into seven functional categories: immune cells, tissue cells, cytokines, immune cell processes, metabolism, T-cell subsets, and skin barrier–related pathways.

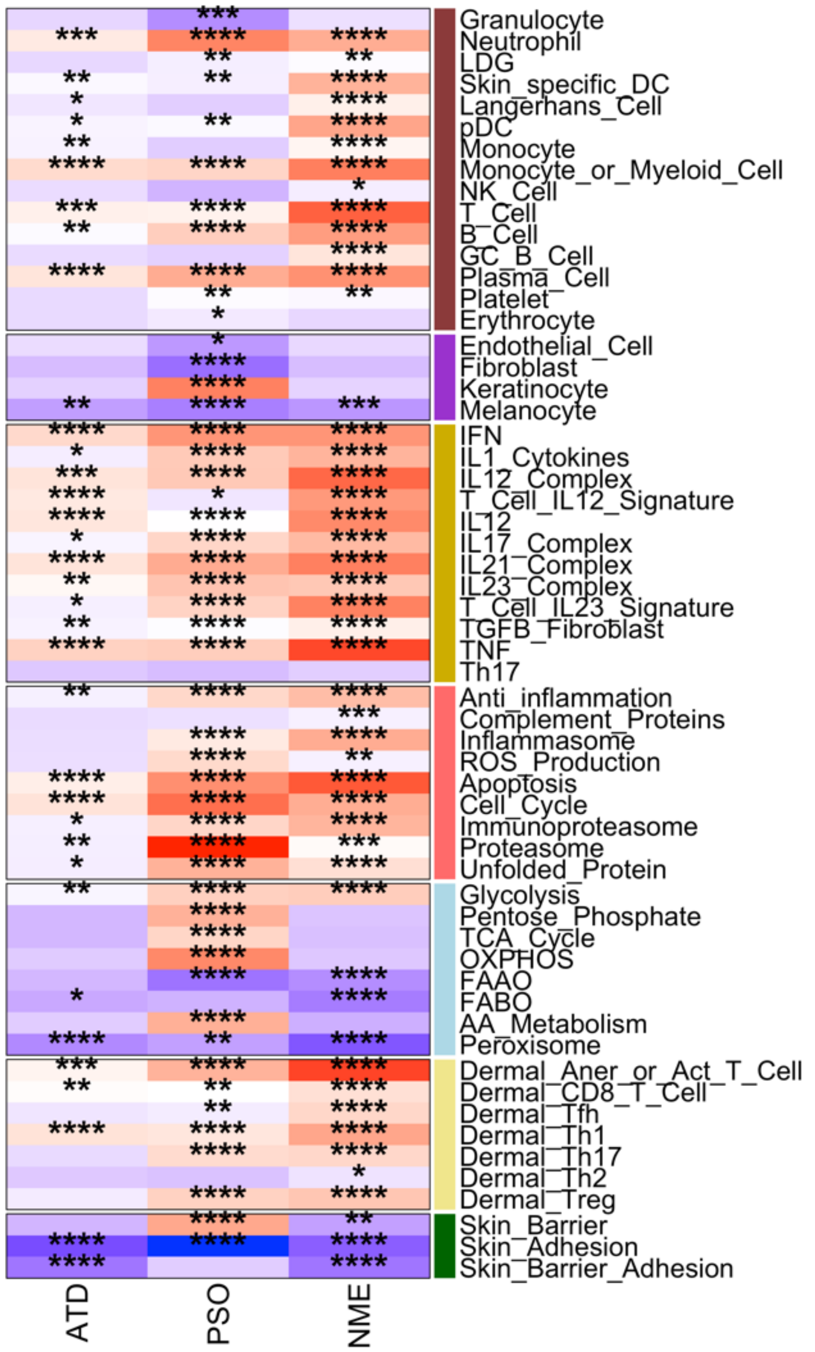


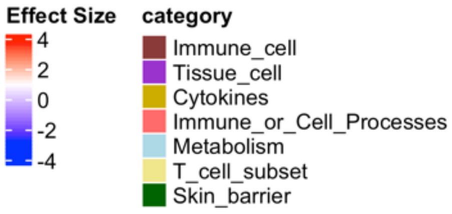


**<u>Figure S5: Transcriptomic comparison of lesional (LES) and non-lesional (NLS) skin in atopic dermatitis (ATD), psoriasis (PSO), and nummular eczema (NME).</u>** Heatmap showing Hedges' g effect sizes for 56 gene modules comparing LES versus NLS skin within each disease. Modules are grouped into seven categories: immune cells, cytokines, tissue cells, immune cell processes, metabolism, T-cell subsets, and skin barrier pathways. Positive effect sizes (red) indicate higher GSVA enrichment in LES skin, while negative values (blue) indicate lower enrichment. Asterisks denote significance from paired t-tests (*P < 0.05; **P < 0.01; ***P < 0.001; ****P < 0.0001). The analysis highlights both shared and disease-specific transcriptomic differences between LES and NLS skin.

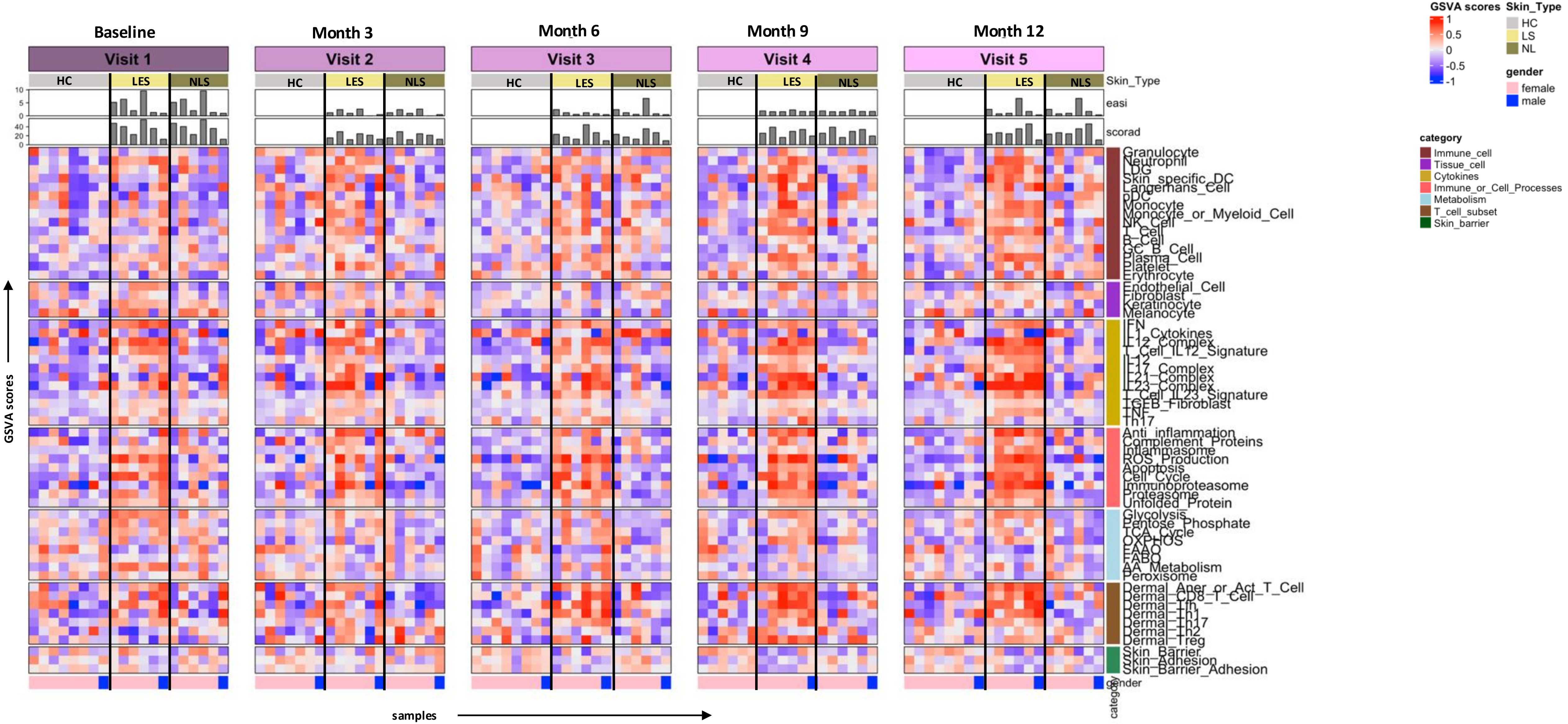


**Figure S6: Dynamic changes in AD pathogenesis over time**: Heatmap representing the GSVA scores for LES and NLSs skin samples from untreated adult AD patients over time, compared to healthy controls. The data spans several timepoints including baseline, month 3, month 6, month 9 and month 12. Additionally, EASI and SCORAD are presented as bar graphs above the heatmap, providing a visual reference for disease severity at each time point.

**Figure S7: Longitudinal changes in immune gene signatures in untreated lesional and non-lesional skin of AD patients and healthy controls**. Violin plots show GSVA scores for selected immune and skin-associated gene modules across lesional (AD_LS), non-lesional (AD_NL), and control (CTL) skin at baseline, month 3, 6, 9, and 12. Modules include T-cell subsets (Dermal_Th1, Dermal_Th2, Dermal_Treg), cytokine signaling (IFN, IL1_Cytokines, IL12), tissue-resident cells (Erythrocyte, Skin-specific-DC), and innate immune processes (Inflammasome). Each violin represents score distribution per timepoint, with dots indicating individual patient GSVA values. Statistical comparisons were performed within each tissue type across timepoints using ANOVA. Asterisks denote significant temporal differences (*P < 0.05; **P < 0.01).

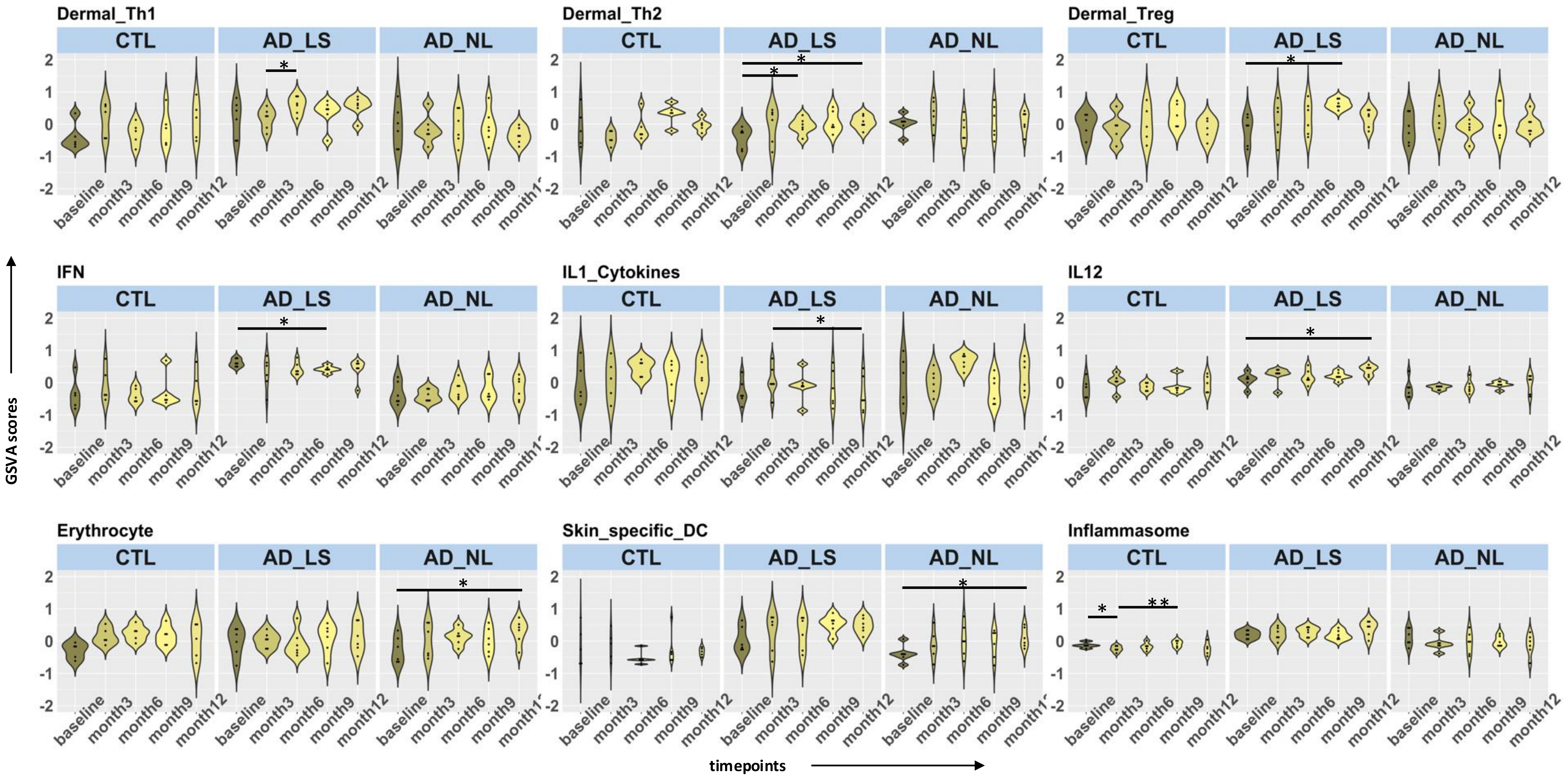

Step 1 Concatenated GSVA scores of baseline LS and healthy samples across multiple datasets

Step 2 Logistic regression with ridge penalty

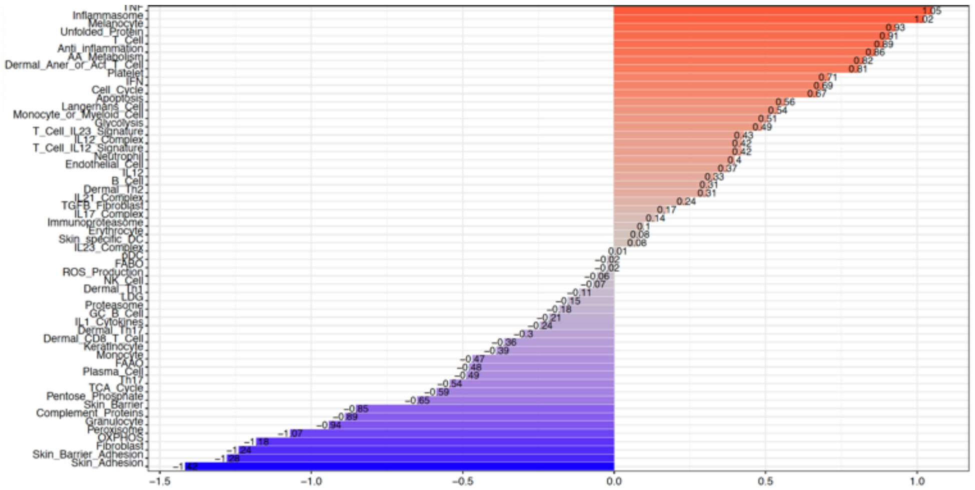


Step 3 GSVA score multiplied by coefficient

Step 4 Normalization by adding minimum score

Step 5 **Eczema Cell and Immune Score (ECZECIS)**

**Figure S8: Derivation and correlation of Eczema Cell and Immune Score (ECZECIS):** Flow diagram illustrating the methodology used to derive the ECZECIS. Coefficients are visualized in bar graph

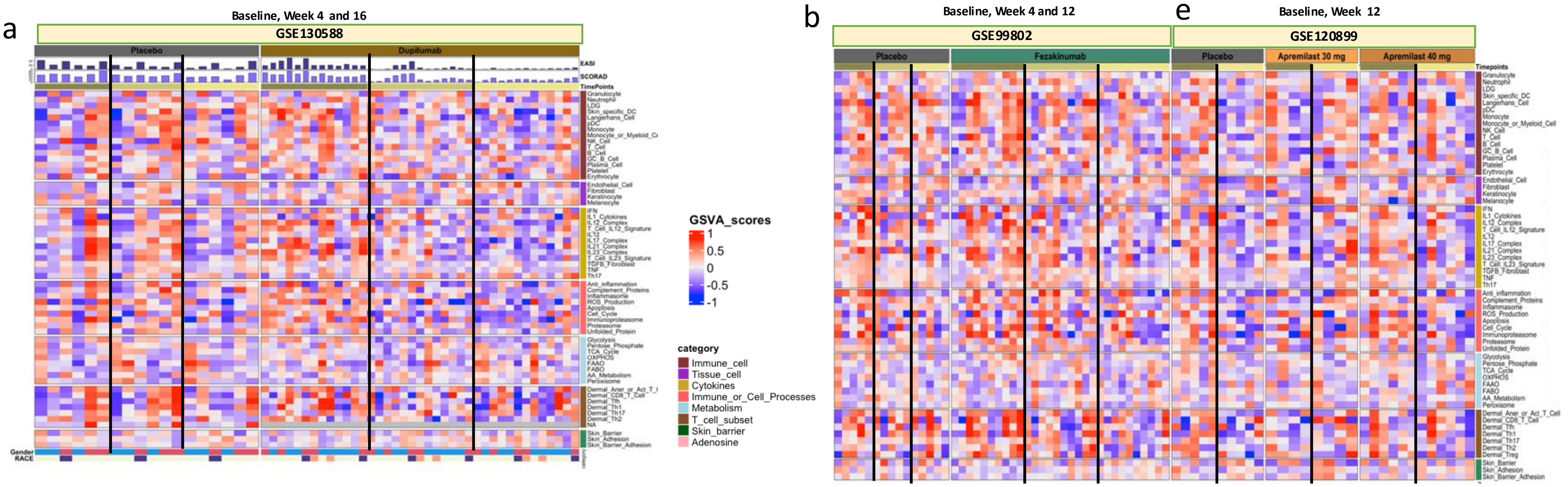


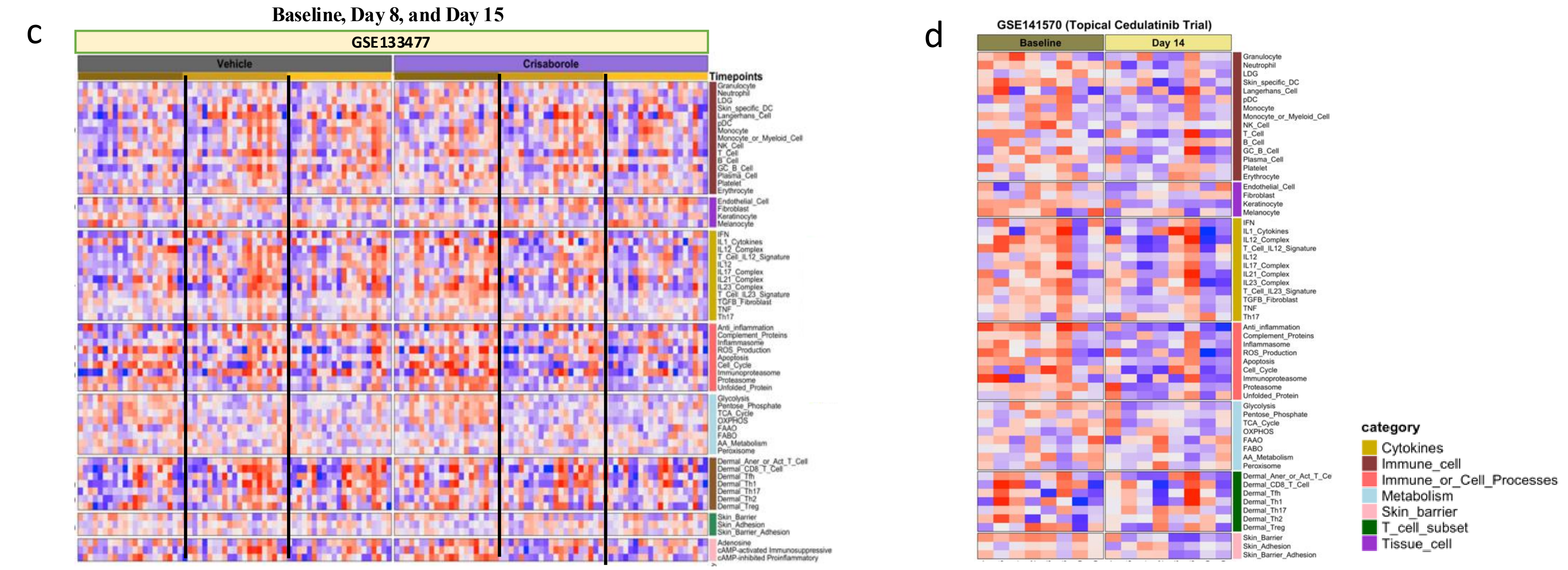


**Figure S9: Longitudinal changes in the gene expression profiles of AD patients treated with various therapeutic agents:** **a-d.** Heatmaps displaying GSVA scores for LS skin samples from AD patients treated with **a**. Placebo or Dupilumab at baseline, week 4, and week 16 (GSE130588). **b.** Placebo or Fezakinumab at BL, Week 4 and Week 12, **c**. Apremilast (30 mg or 40 mg) at baseline, and week 12. **d**. Placebo or Crisaborole at baseline, Day 8, and Day 15. **e**. Cedulatinib at baseline and Day 14

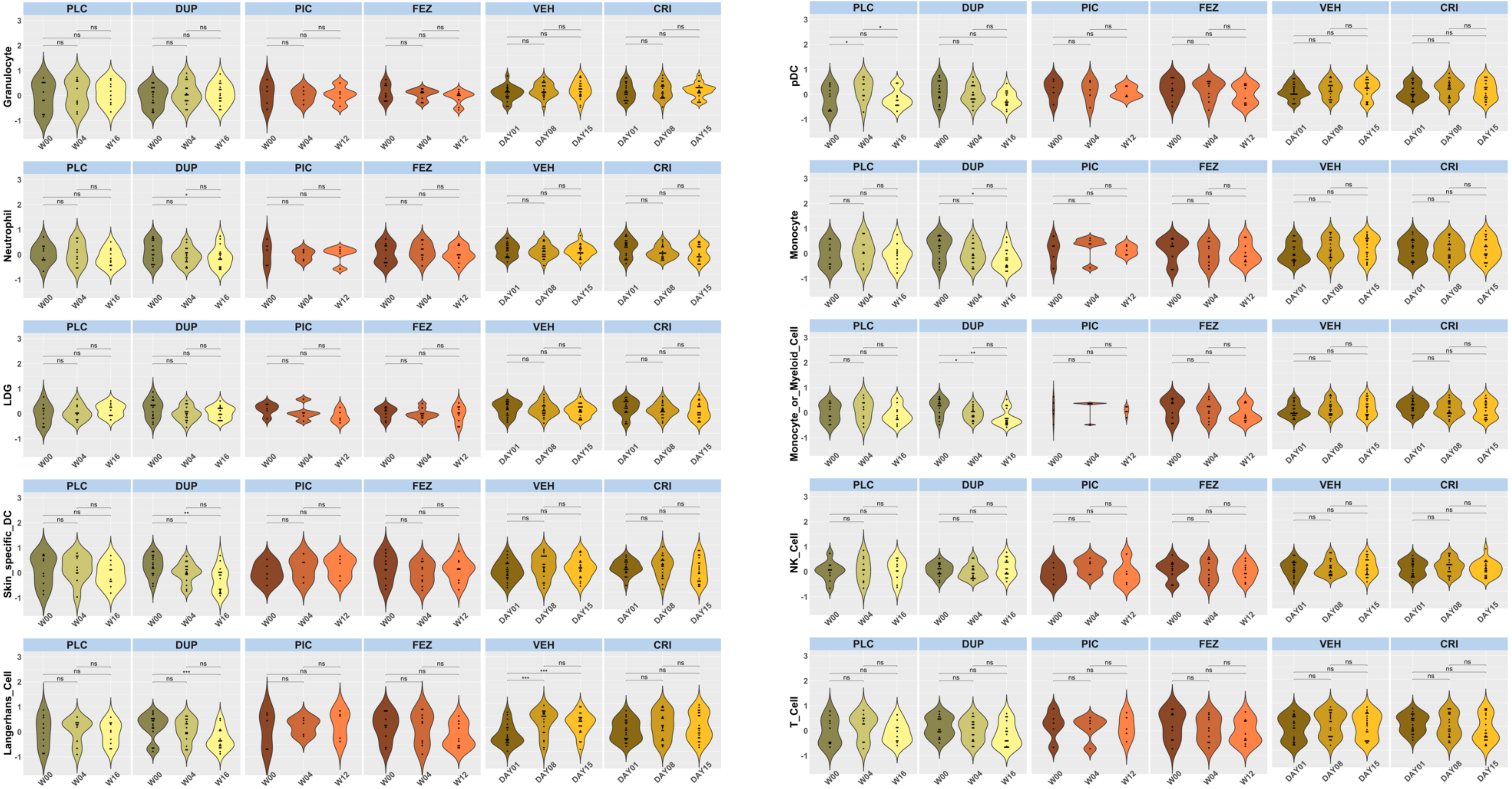
PLC
DUP
PIC
FEZ
VEH
CRI
Granulocyte
Neutrophil
LDG
Skin_specific_DC
Langerhans_Cell
pDC
Monocyte
Monocyte_or_Myeloid_Cell
NK_Cell
T_Cell
W00
W04
W16
W12
DAY01
DAY08
DAY15

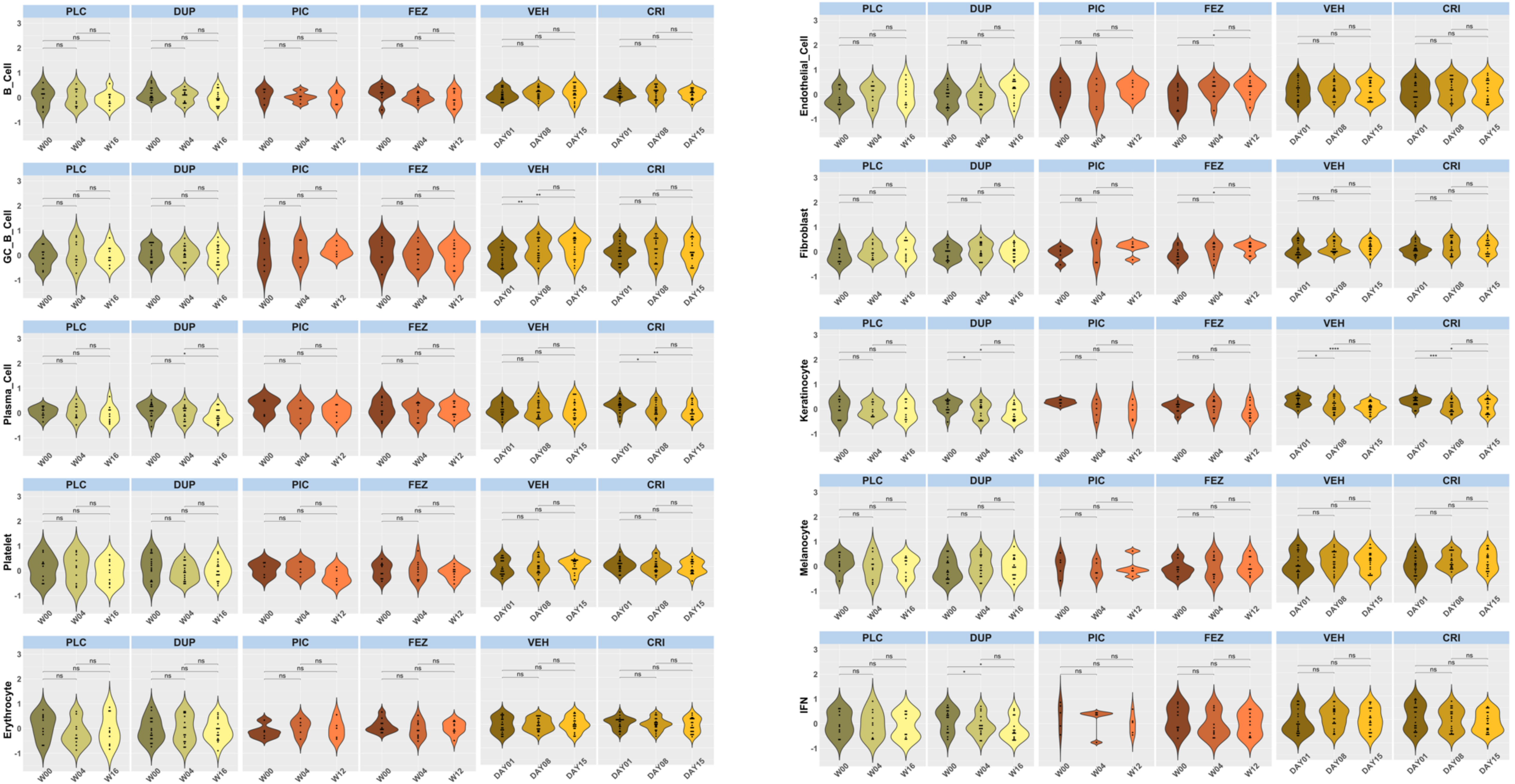

PLC
DUP
PIC
FEZ
VEH
CRI
B_Cell
GC_B_Cell
Plasma_Cell
Platelet
Erythrocyte
Endothelial_Cell
Fibroblast
Keratinocyte
Melanocyte
IFN
W00
W04
W16
W12
DAY01
DAY08
DAY15
ns

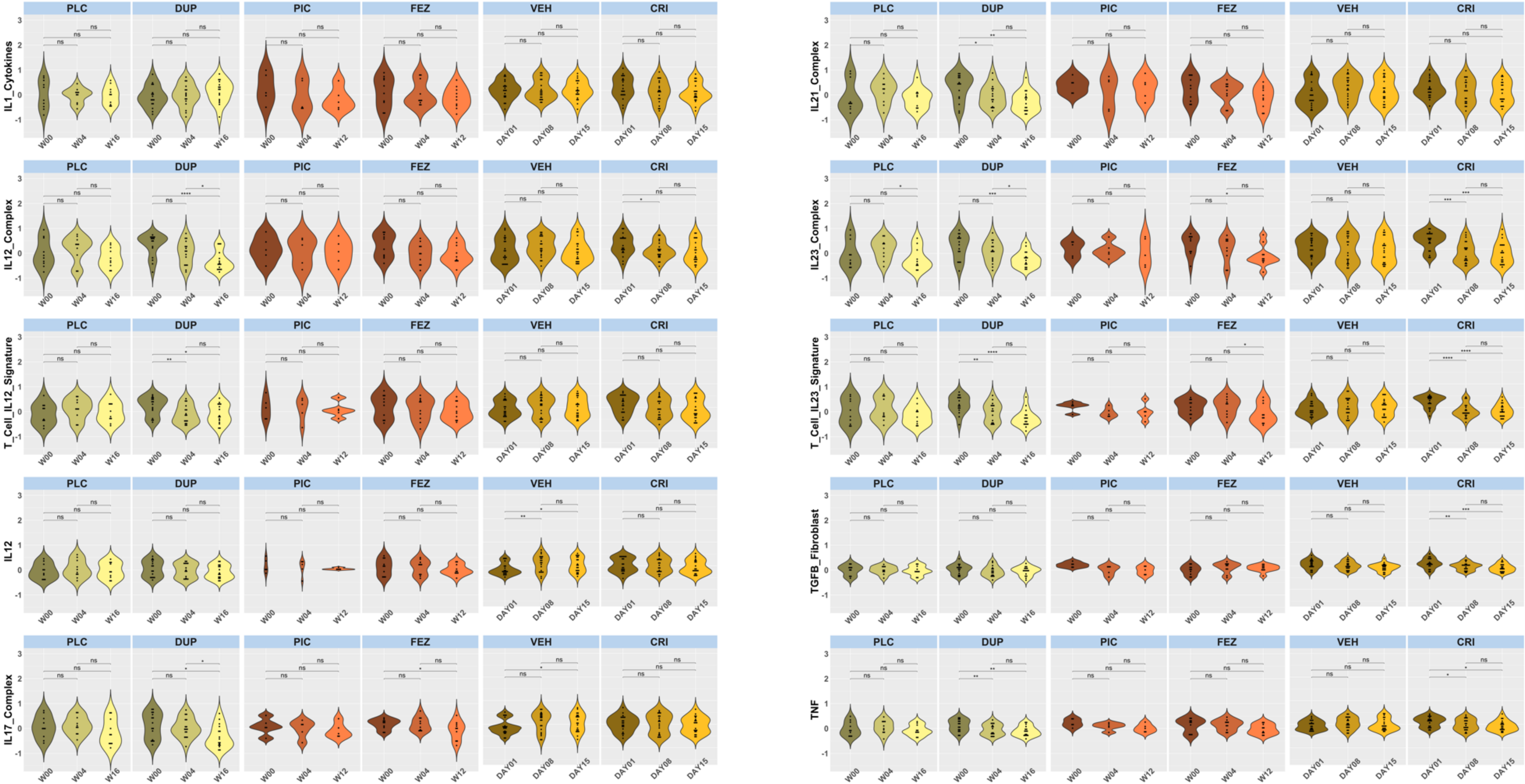
PLC
DUP
PIC
FEZ
VEH
CRI
IL1_Cytokines
IL12_Complex
T_Cell_IL12_Signature
IL12
IL17_Complex
IL21_Complex
IL23_Complex
T_Cell_IL23_Signature
TGFB_Fibroblast
TNF
W00
W04
W16
W12
DAY01
DAY08
DAY15

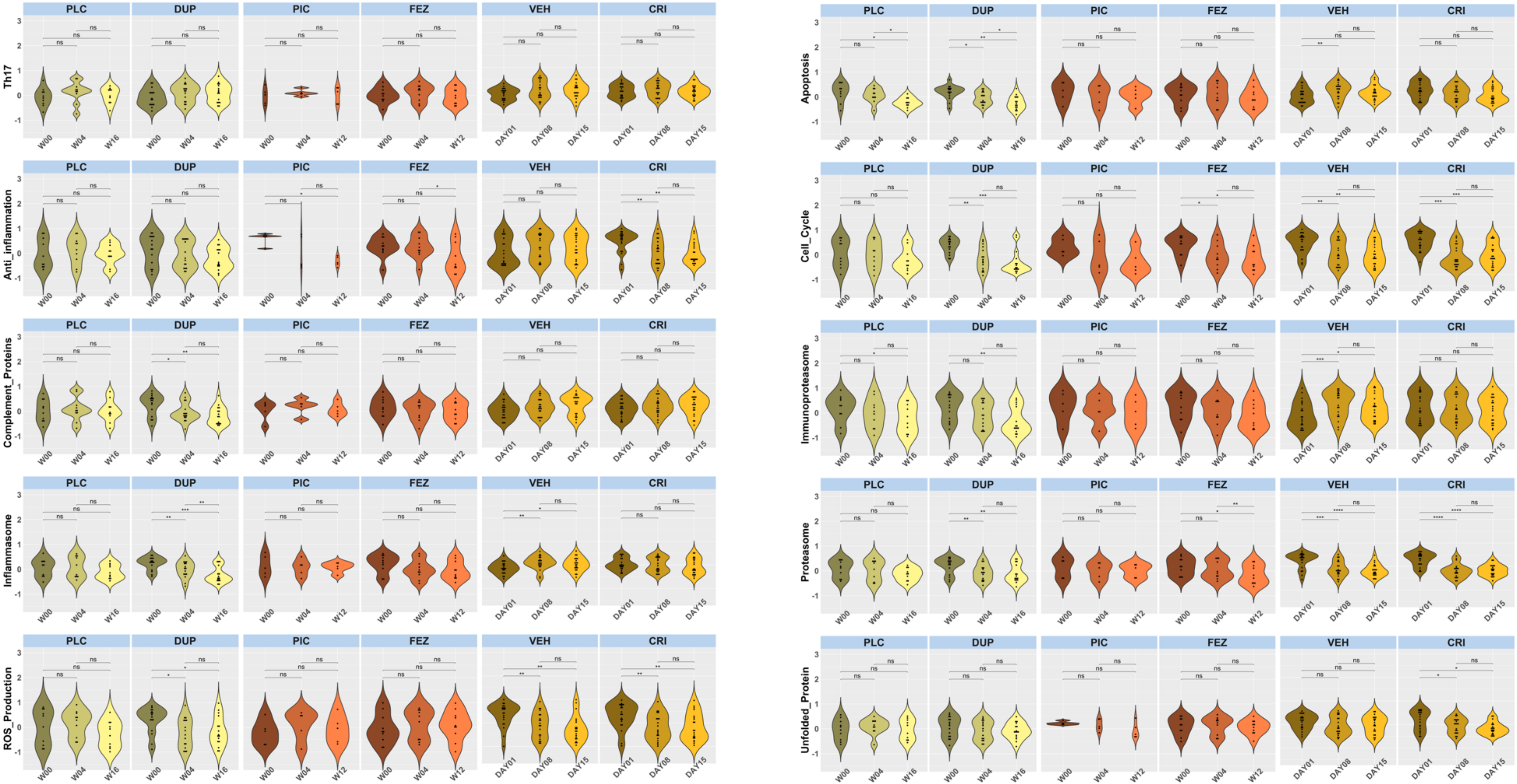
PLC
DUP
PIC
FEZ
VEH
CRI
Th17
Anti_inflammation
Complement_Proteins
Inflammasome
ROS_Production
Apoptosis
Cell_Cycle
Immunoproteasome
Proteasome
Unfolded_Protein
W00
W04
W16
W12
DAY01
DAY08
DAY15
ns

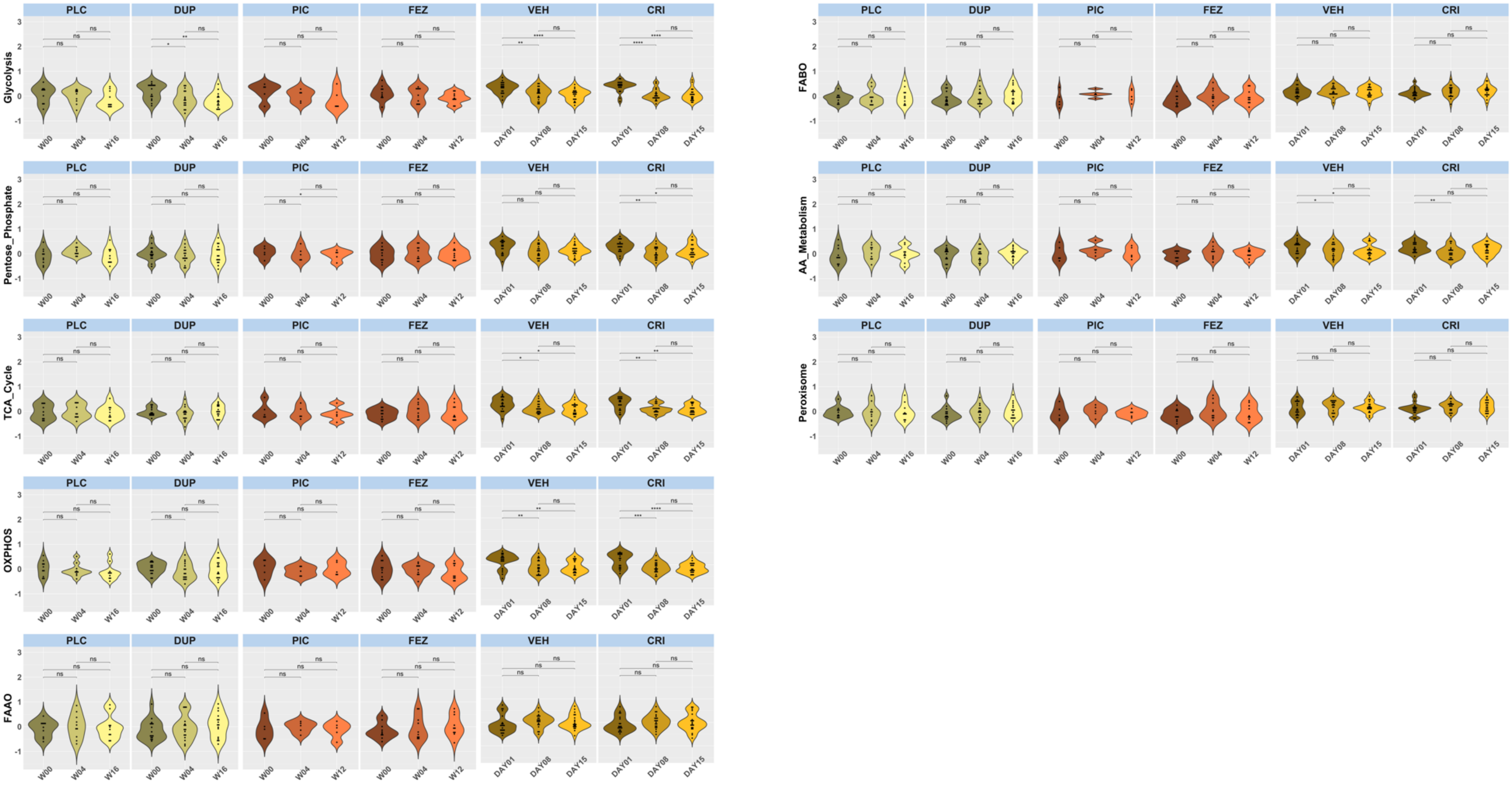

PLC
DUP
PIC
FEZ
VEH
CRI
Glycolysis
Pentose_Phosphate
TCA_Cycle
OXPHOS
FAAO
FABO
AA_Metabolism
Peroxisome
W00
W04
W16
W12
DAY01
DAY08
DAY15
ns
*
**
***
****

### T cell subsets

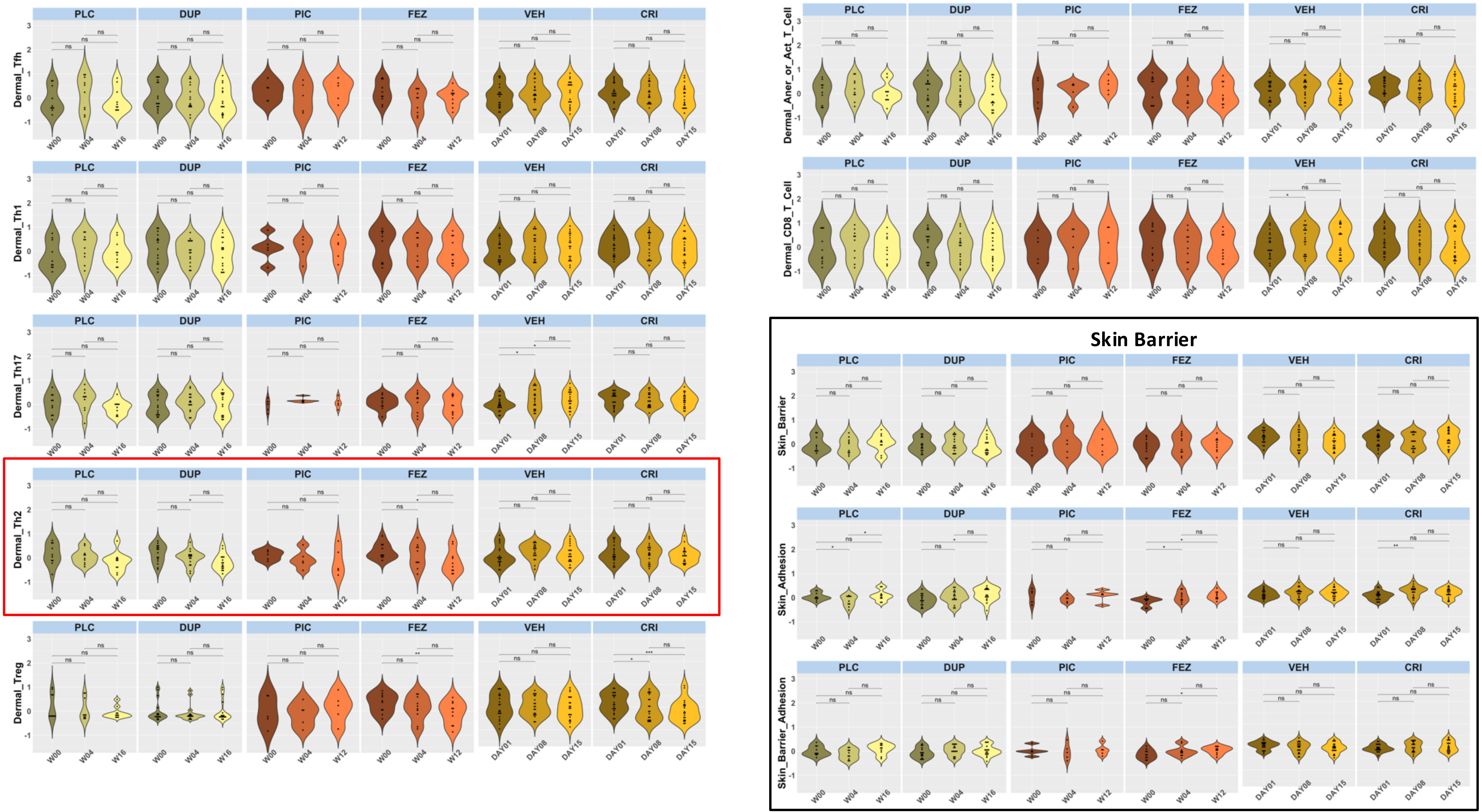

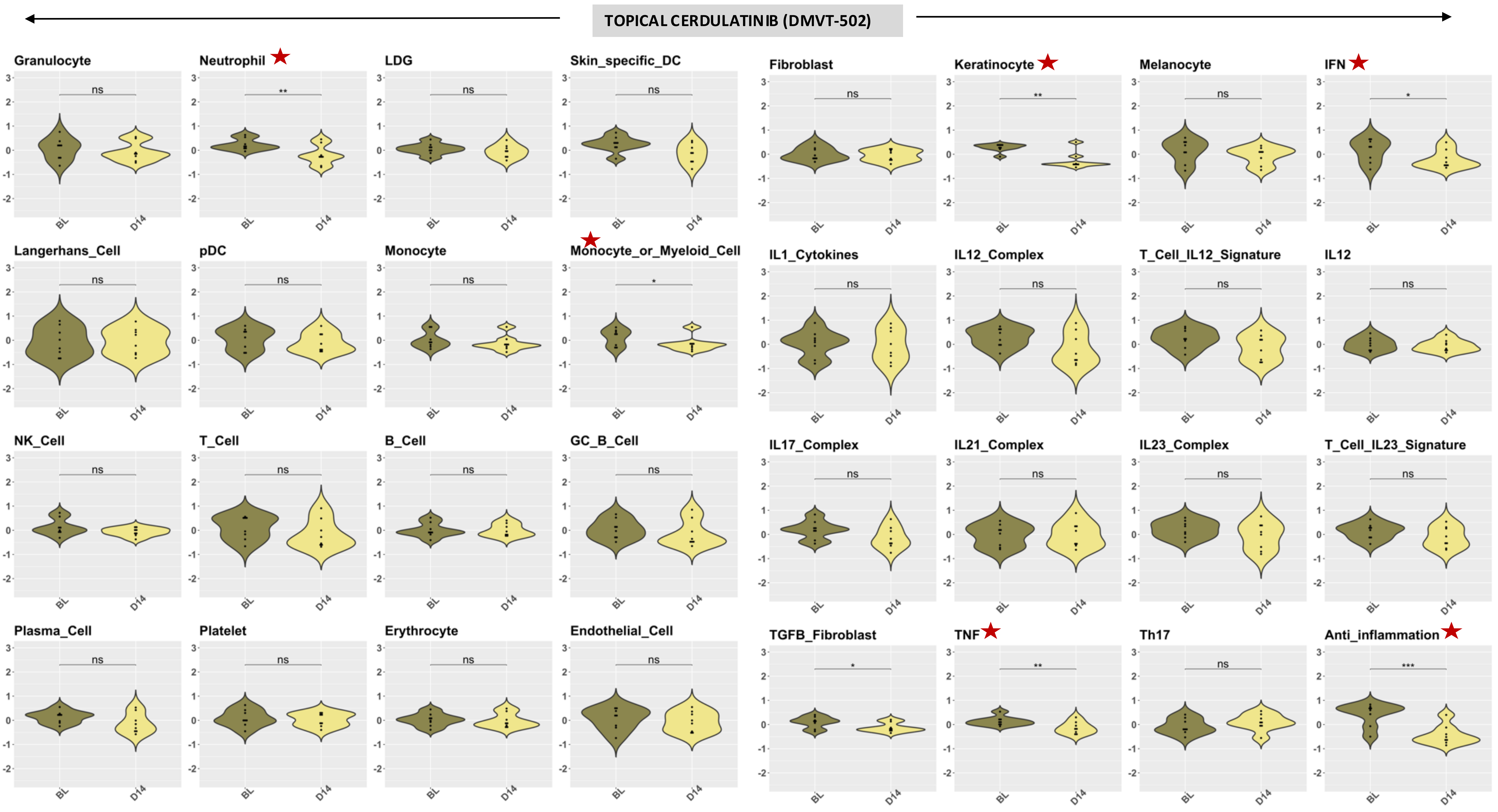
TOPICAL CERDULATINIB (DMVT-502)
Granulocyte
Neutrophil
LDG
Skin_specific_DC
Fibroblast
Keratinocyte
Melanocyte
IFN
Langerhans_Cell
pDC
Monocyte
Monocyte_or_Myeloid_Cell
IL1_Cytokines
IL12_Complex
T_Cell_IL12_Signature
IL12
NK_Cell
T_Cell
B_Cell
GC_B_Cell
IL17_Complex
IL21_Complex
IL23_Complex
T_Cell_IL23_Signature
Plasma_Cell
Platelet
Erythrocyte
Endothelial_Cell
TGFB_Fibroblast
TNF
Th17
Anti_inflammation
BL
D14

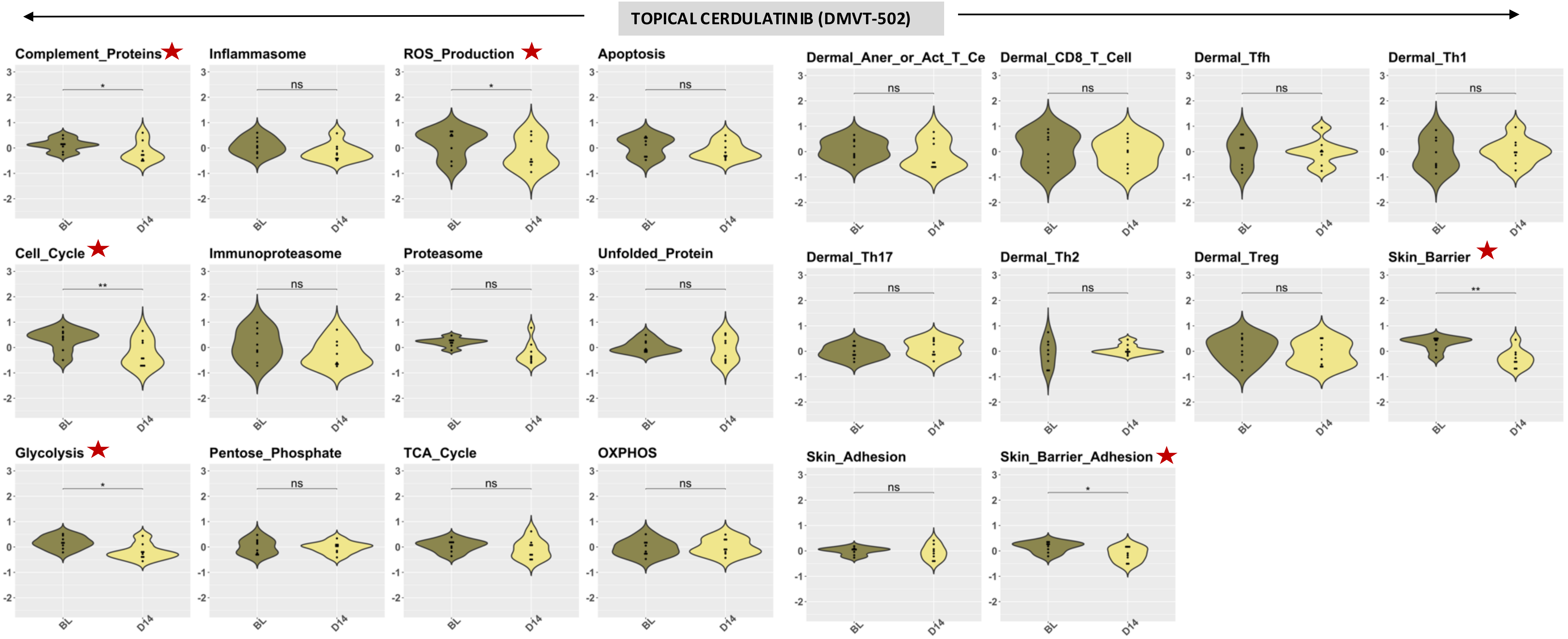


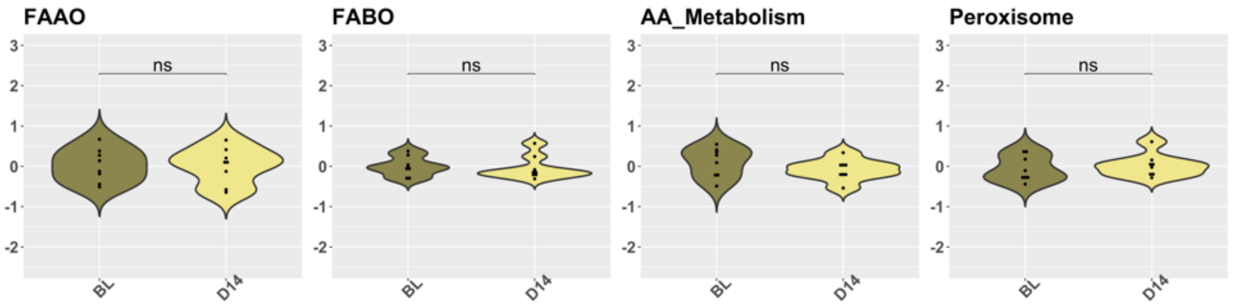


**<u>Figure S10: Longitudinal changes in immune cell gene signatures in lesional skin of AD patients treated with various agents.</u>** Violin plots display GSVA scores for 56 immune-related gene modules in lesional (LES) skin from patients receiving placebo (PLC), dupilumab (DUP), fezakinumab (FEZ), vehicle (VEH), crisaborole (CRI) or Cedulatinib. Scores are shown at baseline (W0), intermediate (W4 for PLC, DUP, FEZ; Day 8 for VEH, CRI), and final timepoints (W12, W16, Day 15, or Day 14). Each violin shows GSVA score distribution per timepoint; dots indicate individual patient values. Statistical comparisons were performed within each treatment group across timepoints using ANOVA with multiple-comparison adjustment. Asterisks denote significance (*P < 0.05; **P < 0.01; ***P < 0.001); ns = not significant.

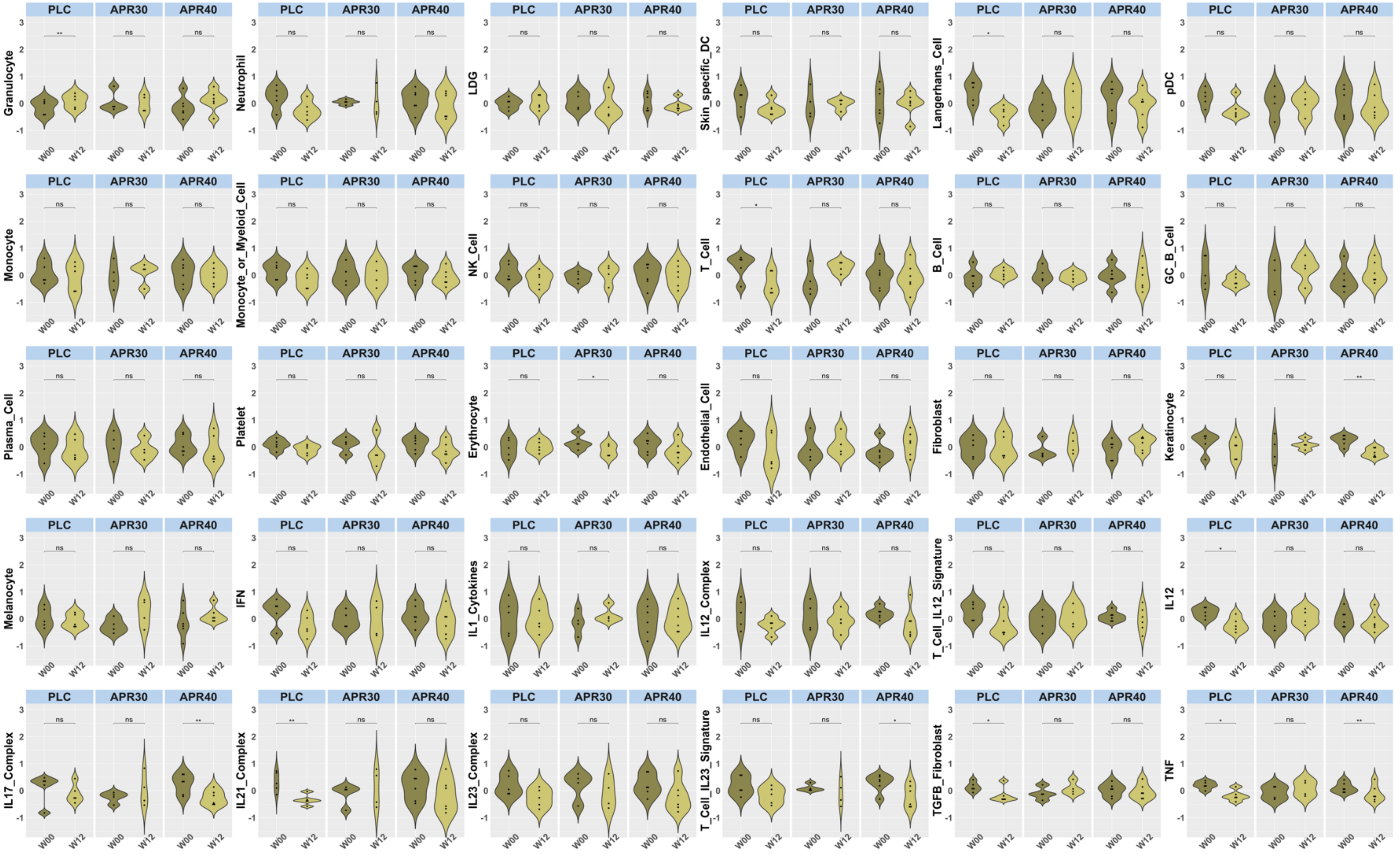
PLC
APR30
APR40
W00
W12
3
2
1
0
-1
Granulocyte ** ns ns
Neutrophil ns ns ns
LDG ns ns ns
Skin_specific_DC ns ns ns
Langerhans_Cell * ns ns
pDC ns ns ns
Monocyte ns ns ns
Monocyte_or_Myeloid_Cell ns ns ns
NK_Cell ns ns ns
T_Cell * ns ns
B_Cell ns ns ns
GC_B_Cell ns ns ns
Plasma_Cell ns ns ns
Platelet ns ns ns
Erythrocyte ns * ns
Endothelial_Cell ns ns ns
Fibroblast ns ns ns
Keratinocyte ns ns **
Melanocyte ns ns ns
IFN ns ns ns
IL1_Cytokines ns ns ns
IL12_Complex ns ns ns
T_Cell_IL12_Signature ns ns ns
IL12 * ns ns
IL17_Complex ns ns **
IL21_Complex ** ns ns
IL23_Complex ns ns ns
T_Cell_IL23_Signature ns ns *
TGFB_Fibroblast * ns ns
TNF * ns **

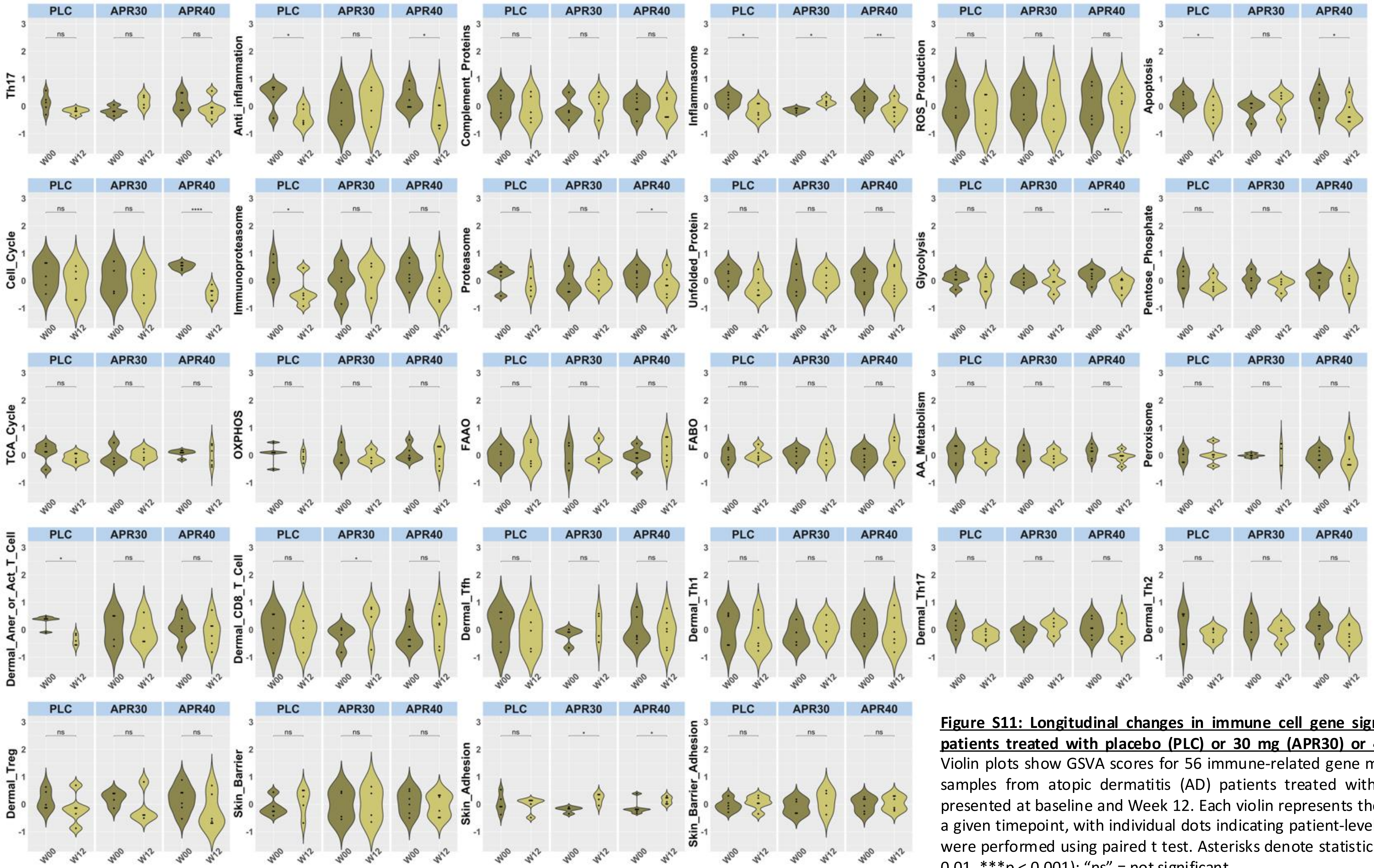


**Figure S11: Longitudinal changes in immune cell gene signatures in lesional skin of AD patients treated with placebo (PLC) or 30 mg (APR30) or 40 mg (APR40) of Apremilast:** Violin plots show GSVA scores for 56 immune-related gene modules across lesional (LS) skin samples from atopic dermatitis (AD) patients treated with Apremilast. GSVA scores are presented at baseline and Week 12. Each violin represents the distribution of GSVA scores at a given timepoint, with individual dots indicating patient-level values. Statistical comparisons were performed using paired t test. Asterisks denote statistical significance (*p < 0.05, **p < 0.01, ***p < 0.001); "ns" = not significant.

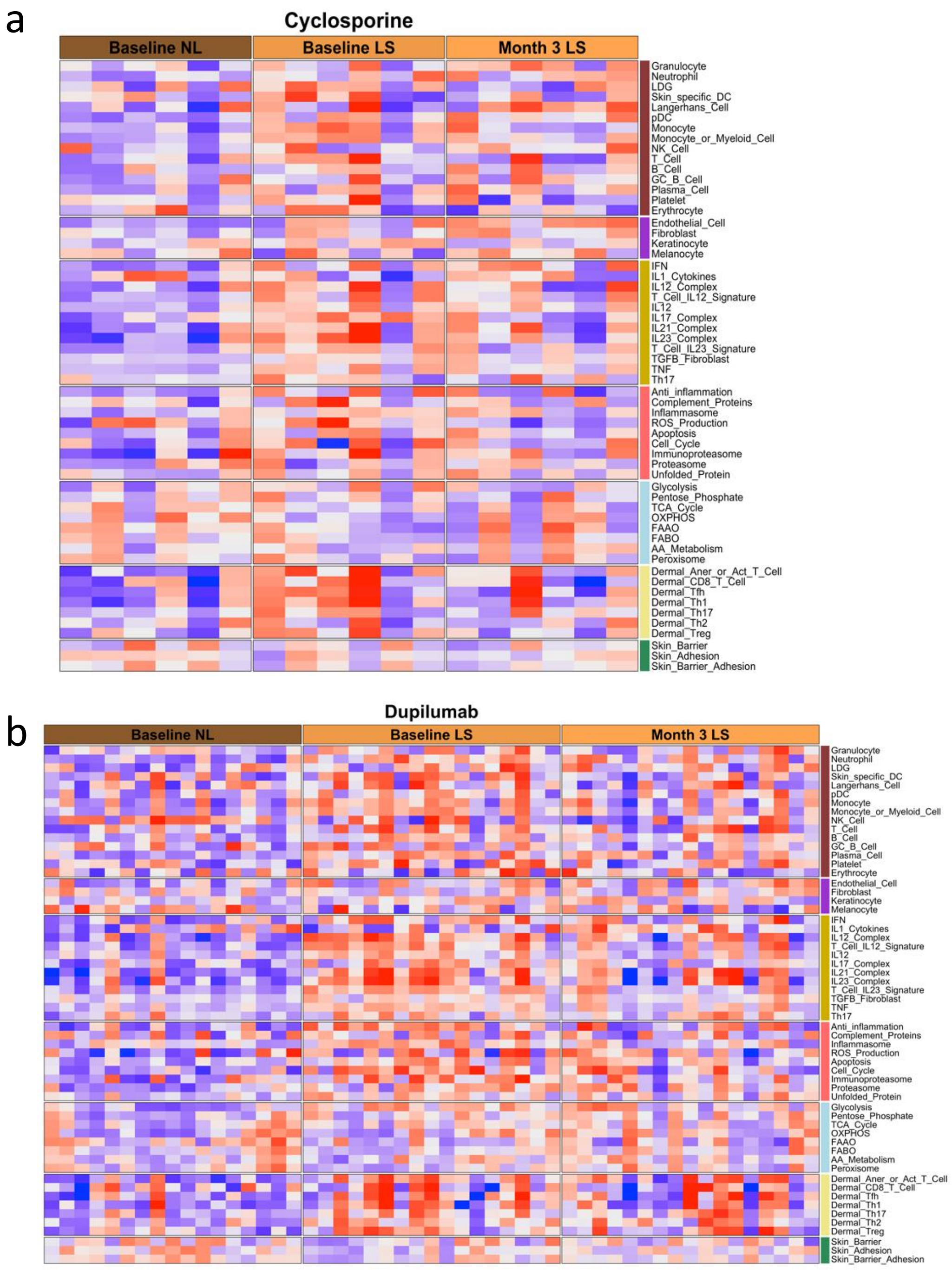


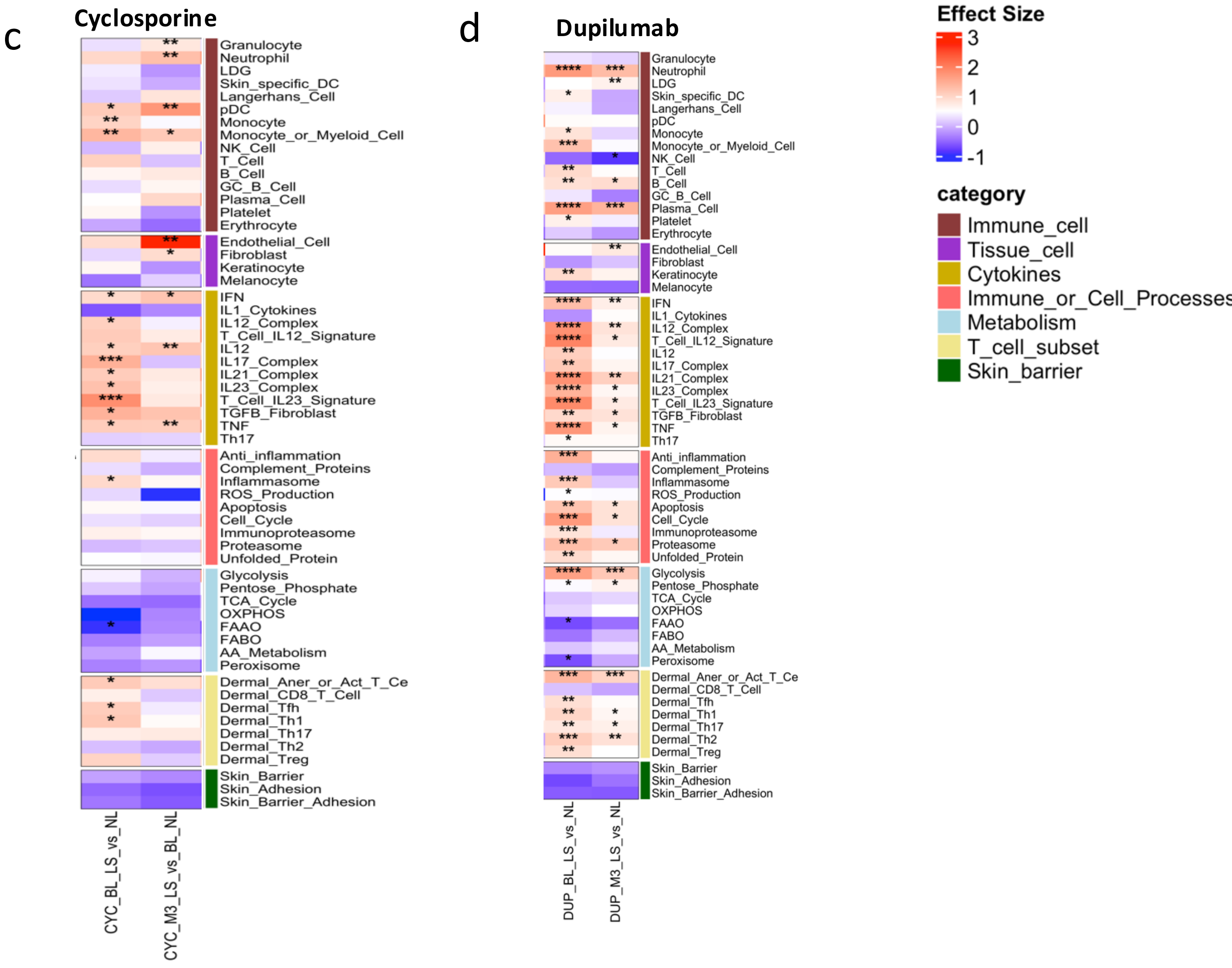


**<u>Figure S12: Transcriptomic impact of cyclosporine and dupilumab treatment in lesional skin of AD patients.</u>** (a-b) Heatmaps showing GSVA scores for 56 curated gene signatures across non-lesional (NLS) and lesional (LES) skin samples of atopic dermatitis (AD) patients treated with **cyclosporine** (a) or **dupilumab** (b), at baseline and Month 3. (c-d) The heatmaps depicts magnitude of change in GSVA scores for each gene signature expressed as Hedge's G effect sizes, across different comparisons: baseline LS versus NL skin, and Month 3 LS vs baseline NL. Red represents an increase in effect size, while blue denotes a decrease. Asterisks indicate statistical significance, determined using paired t-test. Significance levels are denoted by: *P < 0.05, **P < 0.01, ***P < 0.001, and ****P < 0.0001.

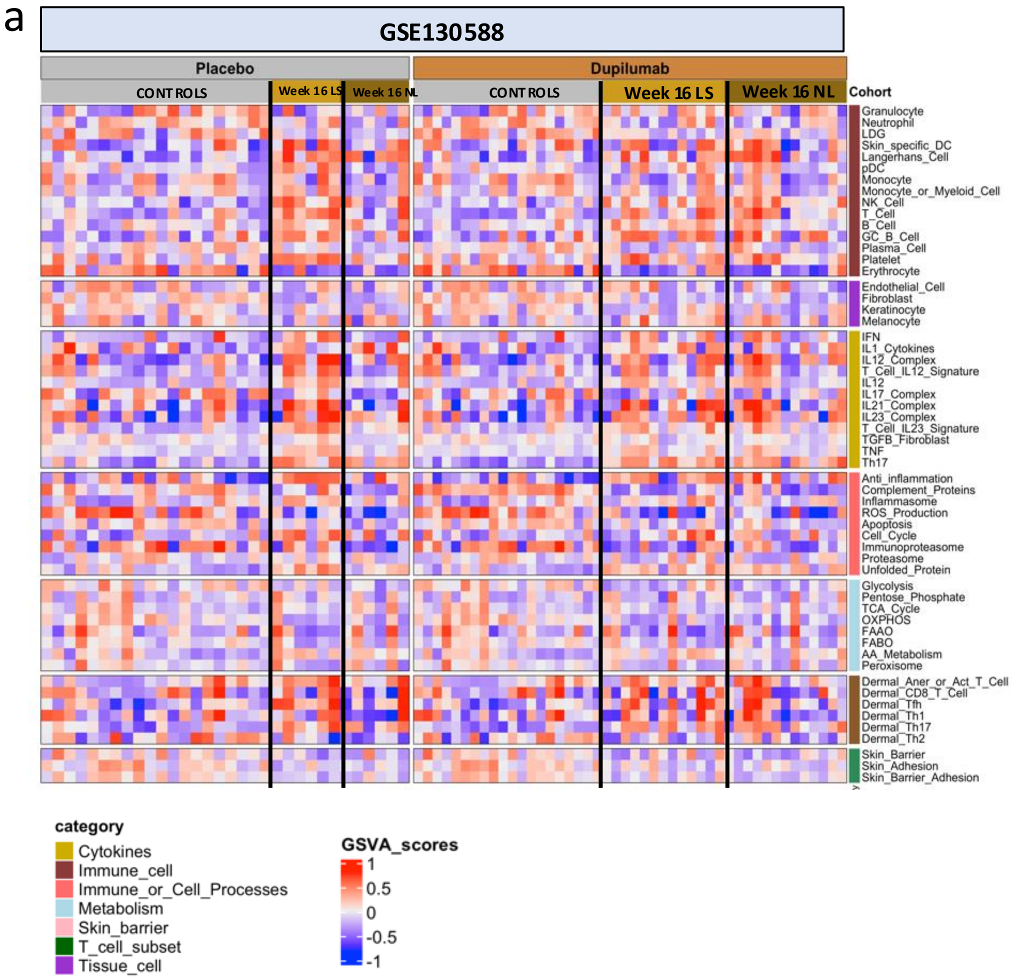


**Figure S13: Transcriptomic comparison of placebo- or dupilumab-treated LES and NLS skin with healthy controls**. **a**. Heatmap showing GSVA scores in AD patients treated with placebo or dupilumab at Week 16, comparing LES and NLS skin with healthy control samples.

a

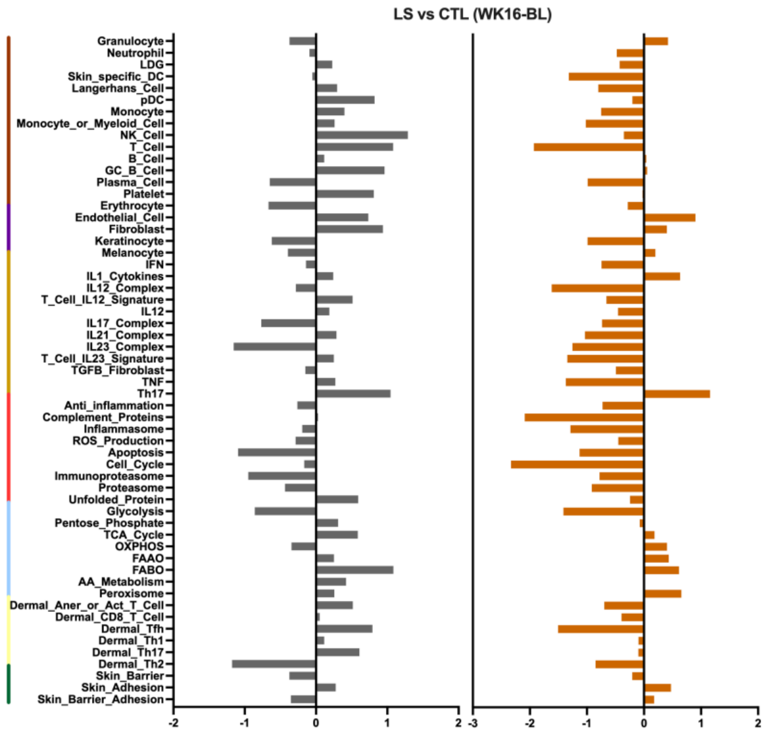


b

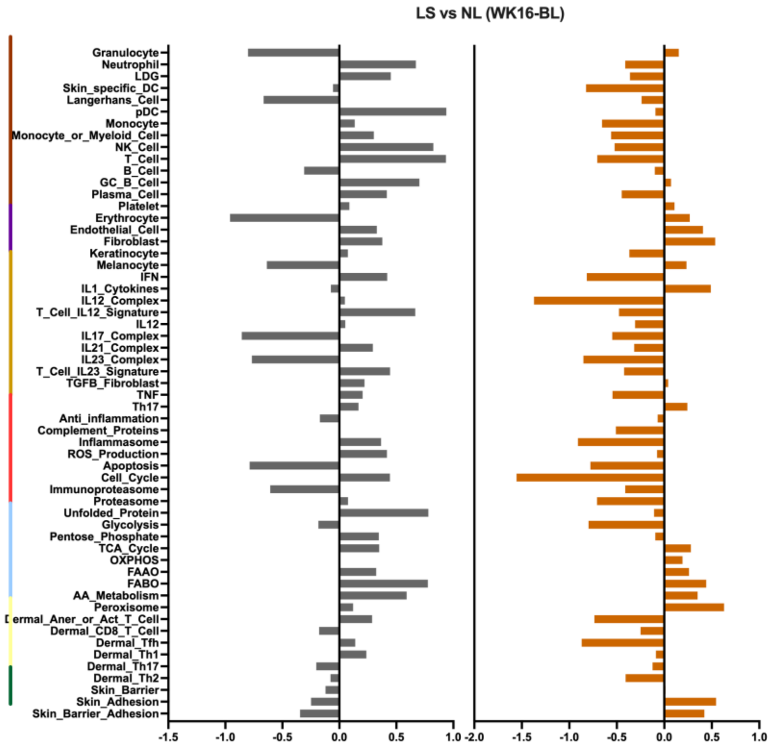


c

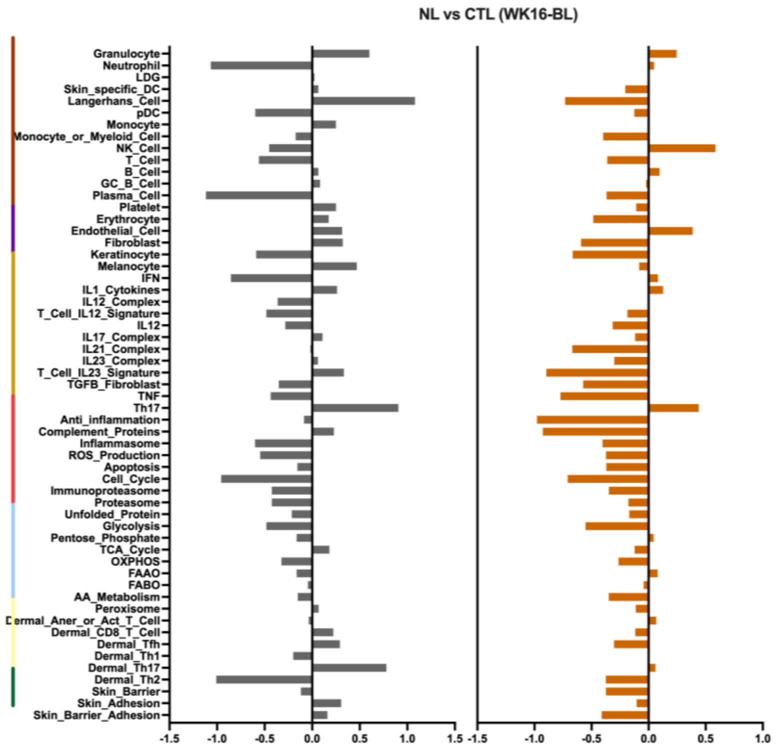


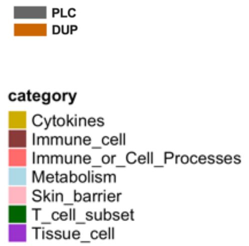


**Figure S14: Transcriptomic comparison of placebo or dupilumab treated LES and NLS skin to healthy controls**: **a-c**. Bar plots showing the change in Hedges' g effect size (Week 16 – Baseline) **a**. LES vs CTL **b**. NLS vs CTL **c.** LES vs NLS skin across multiple gene modules. Separate bars are shown for the placebo and dupilumab treatment groups.
